# A 0.6-meter resolution canopy height model for the contiguous United States

**DOI:** 10.64898/2025.12.12.694075

**Authors:** Scott L. Morford, Brady W. Allred, Shea P. Coons, Anthony A. Marcozzi, Sarah E. McCord, Joseph T. Smith, David E. Naugle

**Affiliations:** Numerical Terradynamic Simulation Group, University of Montana, Missoula, MT, USA; New Mexico Consortium, Los Alamos, NM; Jornada Experimental Range, USDA Agricultural Research Service, Las Cruces, NM, USA; W.A. Franke College of Forestry and Conservation, University of Montana, Missoula MT

## Abstract

Above-ground vertical structure is a critical variable for ecosystem monitoring, carbon accounting, and land management. However, the high cost and limited coverage of airborne lidar hinder its widespread application. To address this, we developed NAIP-CHM, a 0.6-meter resolution canopy height and structure model (CHM) covering the contiguous United States, derived from National Agriculture Imagery Program (NAIP) aerial imagery. Unlike forestry-specific models that exclude human-made features, NAIP-CHM characterizes the full vertical structure of the landscape including vegetation, buildings, and infrastructure. We utilized a U-Net convolutional neural network with attention mechanisms and environmental conditioning, training and validating the model with a peer-reviewed, publicly available dataset of 22.8 million co-registered NAIP imagery and lidar-derived CHM pairs, with stratified sampling to ensure robustness in open-canopy ecosystems. The model achieved a pixel-wise root mean square error (RMSE) of 2.28 meters and an r^2^ of 0.87. Forested sites alone produced an r^2^ of 0.82 and RMSE of 3.82 meters. We provide the dataset, source code, and cloud-based tools to enable broad application without requiring specialized computational resources.

## Background & Summary

Accurate characterization of ecosystem structure is essential for addressing foundational questions in ecology, carbon science, and land management^1–5^. While airborne and terrestrial lidar systems provide the most precise models of above-ground structure, their high acquisition cost constrains large-scale or repeated application. In contrast, public aerial and satellite imagery programs such as the National Agricultural Imagery Program (NAIP), Sentinel-2, and Landsat offer systematic, recurring coverage. When coupled with deep learning, these public resources provide a cost-effective pathway to produce spatially continuous canopy height models (CHMs).

Recent deep learning applications have produced several notable canopy height products. Global-scale mapping at 10-meter resolution has been demonstrated using Sentinel-2 and Global Ecosystem Dynamics Investigation (GEDI) lidar data^6^ and meter-scale global models have been developed from commercial satellite imagery^7^. Regional efforts have further confirmed the utility of high-resolution aerial imagery for fine-scale mapping of canopy structure^8–10^. However, the accessibility of these products is often restricted by proprietary and costly inputs, closed-source code, or specialized expertise required to implement complex models. Furthermore, model development has historically prioritized forested ecosystems, often degrading performance in open-canopy environments like rangelands. Critically, many products mask human-made structures to support carbon accounting, limiting their utility for applications like wildfire modeling and wildlife habitat assessments where the physical presence of both vegetation and infrastructure matters. This represents a significant information gap, as open-canopy ecosystems and the wildland-urban interface are globally significant and central to land management^11,12^.

To address these limitations, we introduce a 0.6-meter resolution canopy height dataset for the contiguous United States (CONUS). Functionally acting as a normalized Digital Surface Model (nDSM) by capturing all elevated features, the dataset is designated NAIP-CHM to highlight its focus on ecosystem structure. We generated the dataset using NAIP imagery spanning 2012 to 2023, with 96% of the imagery acquired in 2022–2023. We employed a U-Net^13^ convolutional neural network with a Convolutional Block Attention Module^14^ (CBAM) to refine feature maps and Feature-wise Linear Modulation^15^ (FiLM) to integrate auxiliary environmental data. The model was trained using a previous published public dataset of over 22 million NAIP and lidar derived CHM pairs^16^.

We prioritize open access by providing the dataset and model through multiple channels suited to different user needs. For free bulk access independent of commercial cloud platforms, Cloud Optimized GeoTIFFS (COGs) are available for direct download from our servers. For users preferring cloud-based workflows, the data are also accessible via Google Earth Engine and a Google Cloud Storage bucket. To facilitate reproducibility and downstream application, we release the complete modeling pipeline, including the trained model, inference scripts, and a Google Colab notebook for applying the model to past and future NAIP imagery. Additionally, an interactive Google Earth Engine application is provided for no-code data exploration. These combined resources ensure the NAIP-CHM dataset is accessible to users with diverse technical expertise and computational capacity.

## Methods

### Model Training Data: NAIP Imagery and Lidar-Derived Canopy Heights

The model was trained using a publicly available, peer-reviewed dataset of co-registered NAIP imagery and lidar-derived canopy height models (CHMs), comprising over 22 million CHM-NAIP pairs across CONUS^16^. The dataset is stratified by EPA Level III ecoregions^17^ and National Land Cover Database^18^ (NLCD) classes.

The NAIP imagery is 4-band (R, G, B, NIR) orthoimagery at 0.6 or 0.3-meter ground sampling distance, collected by states at varying intervals (typically every 2 to 3 years) during the growing season – generally corresponding to ‘leaf on’ conditions. Reference CHMs were generated from USGS 3D Elevation Program^19^ lidar point clouds meeting quality level 2 (QL2) or higher standards, using a standard pit-free algorithm^20^ at 1.0-meter resolution. NAIP and CHM data are co-registered and resampled to a common 432×432-pixel grid in a single reprojection step. This grid covers a fixed 256 × 256 m physical footprint, giving an effective grid spacing of ∼0.59 m (reported throughout as the nominal 0.6 m resolution). The 432-pixel dimension was chosen for compatibility with the U-Net architecture, which requires input dimensions divisible by 16 due to four successive downsampling stages. Predictions are resampled to 0.6 m resolution for the final product. Each NAIP image was temporally matched to its paired lidar acquisition within a four-year window centered on the lidar collection date; the median temporal offset between lidar and NAIP acquisitions is 200 days (See Fig. 2D in Ref. 16). Lidar data span 2014 through 2023 across CONUS, with geographic coverage determined by 3DEP^21^ availability (Fig. 1 in Ref. 16). Following the sampling stratification described in the companion dataset^16^, grassland and shrubland classes were oversampled by a factor of four and pasture by a factor of two. This intentional oversampling improves model performance by increasing exposure to open-canopy rangeland ecosystems, where existing products have historically struggled.

**Figure 1:**
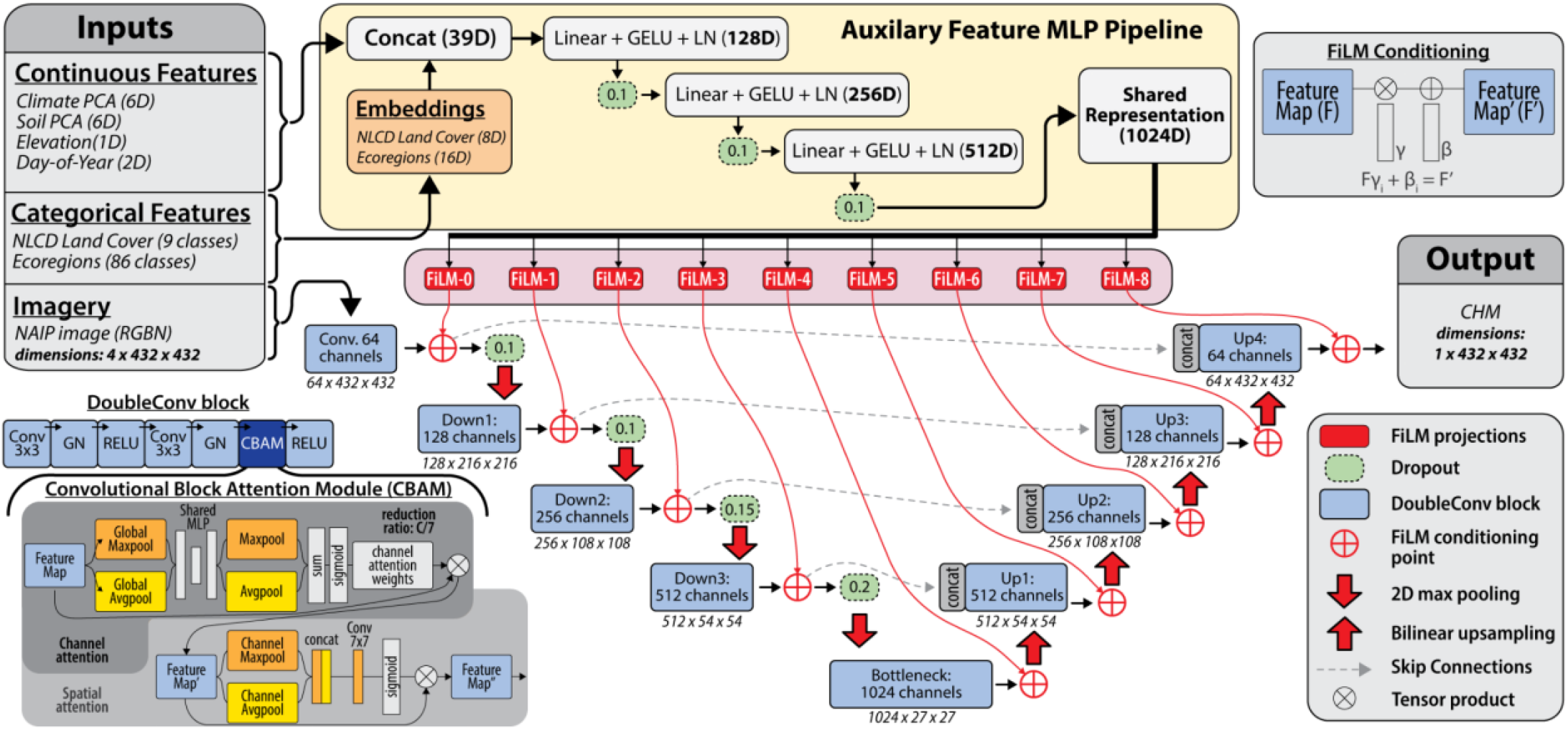
Schematic overview of the architecture used for Canopy Height Model (CHM) prediction. The network fuses high-resolution NAIP imagery with auxiliary environmental variables (e.g., climate, soil, land cover, ecoregion). The auxiliary features are processed via an MLP pipeline to generate a global representation, which dynamically modulates a U-Net backbone via Feature-wise Linear Modulation (FiLM) layers. The U-Net encoder-decoder path is further enhanced with Convolutional Block Attention Modules (CBAM) to refine spatial and channel-wise feature extraction.

**Figure 2:**
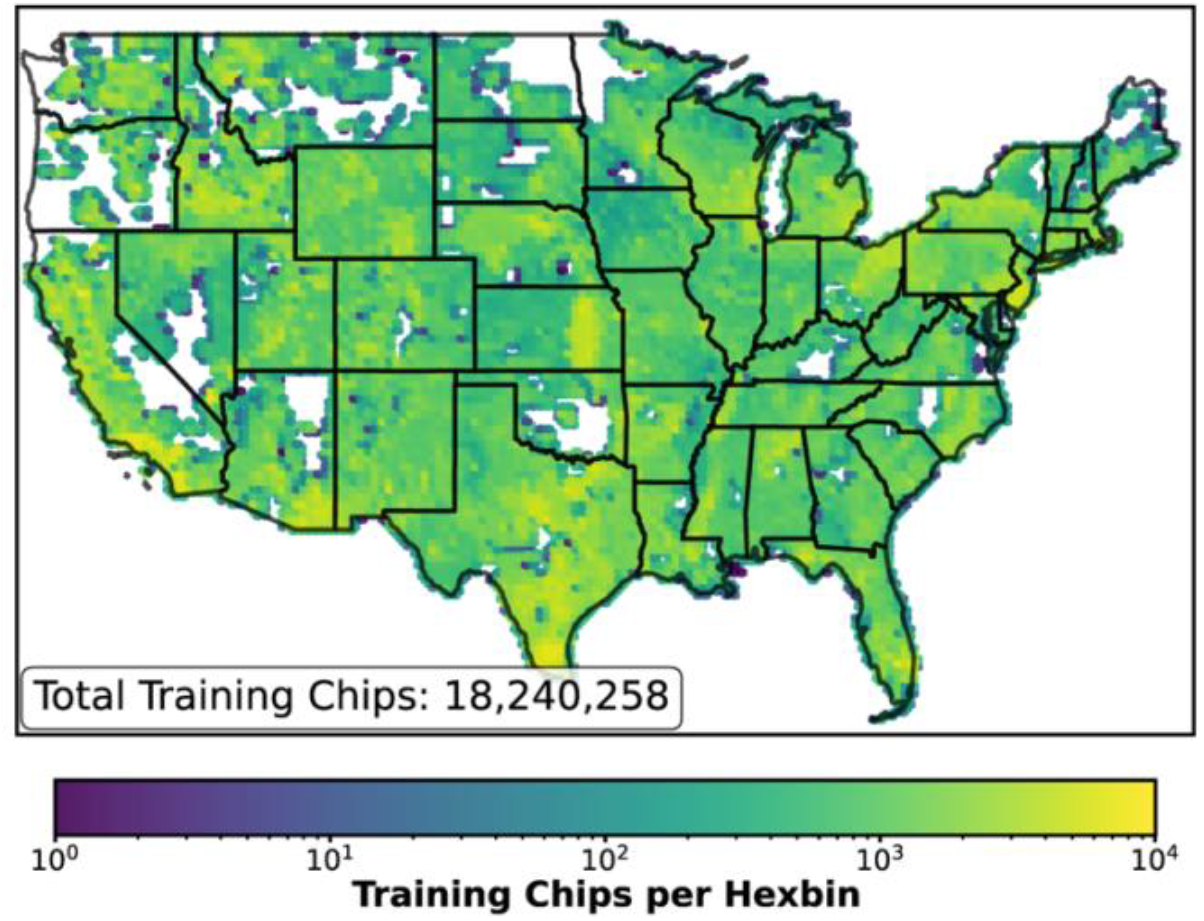
Geographic distribution of the paired NAIP and lidar derived canopy height model (CHM) training dataset. The map displays the spatial density of the 18,240,258 training chips (training partition only) used to develop the model. Hexbin colors represent the count of training samples per area on a logarithmic scale, covering the majority of the contiguous United States.

### Auxiliary Training Data

Auxiliary data were integrated to provide environmental context for model conditioning (**Table 1**). Climate variables were derived from a Principal Component Analysis (PCA) of 20 biologically-relevant features from 30-year PRISM^22^ climatological normals (1991-2020); the first six principal components, capturing over 90% of the variance, were used to account for the influence of temperature and precipitation regimes on vegetation height and species composition. Soil properties were similarly summarized using the first six principal components from a PCA of over 20 variables from the SOLUS database^23^, including texture, bulk density, and soil organic carbon, which impact site productivity through regulation of water availability and nutrient supply. Topography was represented by a digital elevation model from the Shuttle Radar Topography Mission^24^ to capture elevation-driven gradients in temperature, moisture, and species assemblage. The day-of-year of each NAIP acquisition was encoded as sine and cosine components to account for seasonal variation in canopy phenology. Categorical context was provided by a simplified nine-class version of the 2021 NLCD^18^ and EPA Level III ecoregion codes^17^, enabling the model to learn land-cover-specific and regionally varying relationships between spectral inputs and canopy height. In total, 15 continuous and 2 categorical variables were used for model conditioning.

**Table 1:**
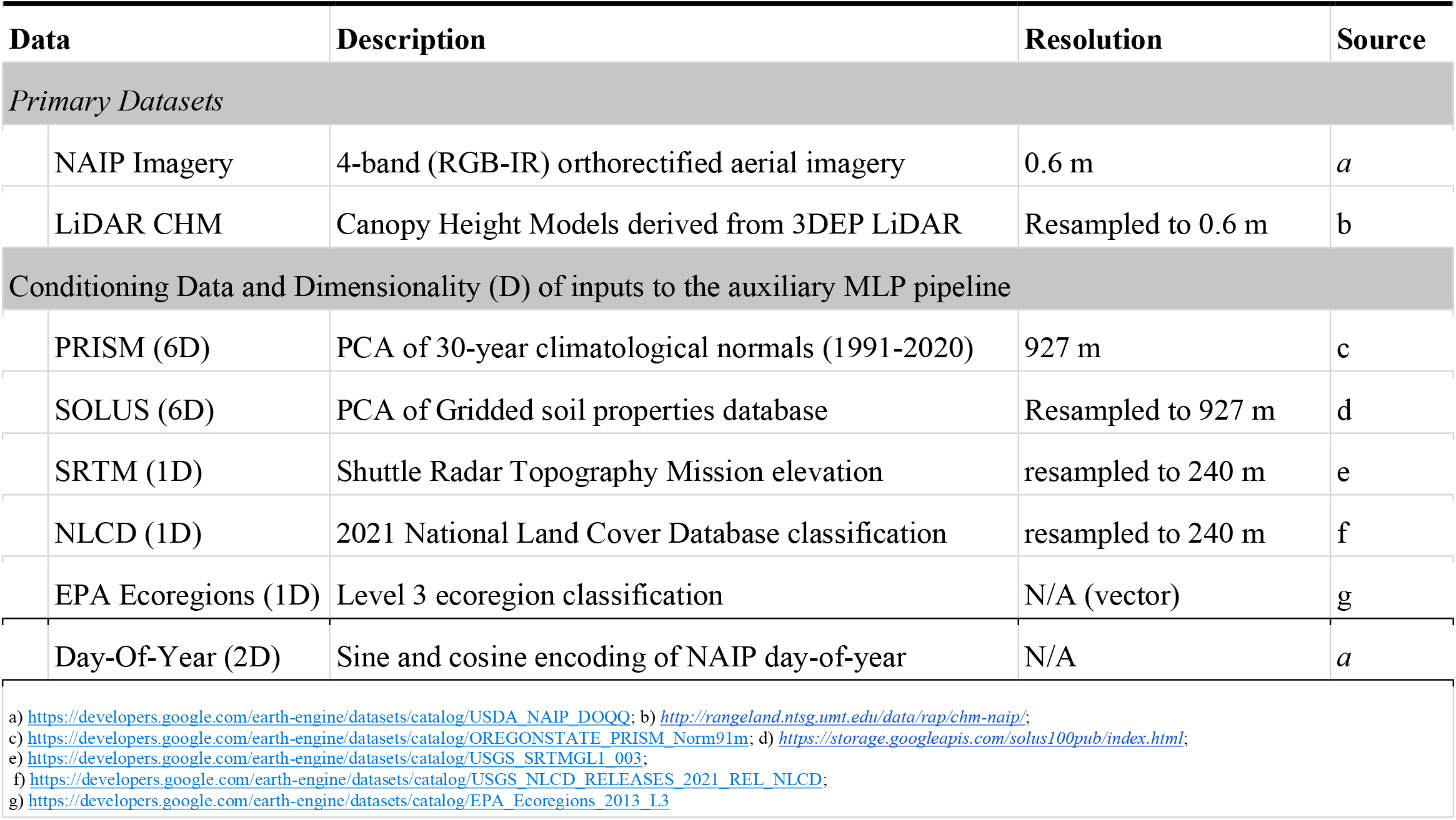
Datasets and Source Repositories Used For the NAIP-CHM Model Development.

### Model Architecture

The model is a U-Net^13^ convolutional neural network (Fig. 1), chosen for its well-established performance in dense remote sensing tasks like canopy mapping^8,10^. It is enhanced with attention mechanisms^14^, which have been shown to improve U-Net accuracy and efficiency^25^. Additionally, multi-stage FiLM^15^ conditioning integrates environmental covariates (climate, soil, elevation, and phenological variables) that modulate learned feature representations at multiple network depths, enabling the model to resolve spectral ambiguities (e.g., distinguishing low-canopy forest from dense shrubland) by conditioning predictions on ecological context rather than image features alone. The model has 22,205,019 parameters and processes two data streams: the 4-band NAIP imagery and the auxiliary environmental features.

The U-Net backbone uses an encoder-decoder structure with channel dimensions of [64, 128, 256, 512, 1024] and skip connections. Each convolutional block consists of two 3×3 convolutions with stride 1 and padding 1, followed by Group Normalization (32 groups) and ReLU activation. These blocks are augmented with CBAM^14^ that applies sequential channel and spatial attention. Downsampling between encoder stages is performed via 2×2 max pooling with stride 2, while upsampling in the decoder uses bilinear interpolation with a scale factor of 2. The encoder employs progressive spatial dropout (rates: 0.1 to 0.2) for regularization. The final output is generated by a 1×1 convolution that maps the decoded features to a single-channel canopy height map.

Auxiliary features are processed in a separate pathway. Categorical features (land cover, ecoregion) are converted to learned embeddings (8 and 16 dimensions, respectively). These are concatenated with the 15 continuous features (climate, soil, elevation, temporal). This 39-dimensional vector is processed by a three-layer multilayer perceptron employing GELU activation and Layer Normalization to produce a 1024-dimensional shared representation.

The FiLM conditioning mechanism modulates U-Net feature maps at nine locations (after the initial convolution and after each encoder and decoder stage). At each location, the shared auxiliary representation is projected to generate channel-specific scale (γ) and shift (β) parameters, which apply affine transformations to the feature maps. This multi-stage conditioning allows the model to integrate environmental information across multiple spatial scales.

### Model Training

Data were partitioned for training, validation, and test sets following the splits provided with the source dataset^16^; the training partition consisted of 18,240,258 paired CHM & NAIP chips (Fig. 2). The model’s inference domain (CONUS) is coextensive with its training domain; random tile-level partitioning therefore matches the geometry of intended deployment ^26–28^.

Training followed a three-phase transfer learning strategy:

1. A U-Net with CBAM was trained from scratch on the 4-band NAIP imagery.
2. The U-Net and CBAM weights were frozen, and only the auxiliary multilayer perceptron and FiLM projection layers were trained.
3. All model weights were unfrozen for end-to-end fine-tuning.

A Huber loss function (delta=1.0) and the AdamW optimizer (learning rate: 6.0e-5, weight decay: 1.0e-4) were used. The learning rate was managed by a ReduceLROnPlateau scheduler. The model was implemented in PyTorch (v2.8.0) and trained on paired NVIDIA H100 GPUs with mixed-precision optimization over approximately four weeks. Geospatial processing used Rasterio (v1.3.0) and PyProj (v3.6.0).

### Model Evaluation

#### Test Dataset

Performance was evaluated on an independent test set with no spatial overlap with training tiles. From the initial test partition of 2,277,602 tiles, a negligible fraction (n = 4,221, 0.19%) was excluded prior to evaluation to remove severe sensor errors or data quality artifacts, resulting in a final evaluation set of n = 2,273,381 tiles. Specifically, we filtered out tiles with insufficient spatial coverage (<25% valid NAIP pixels; n = 3,446, 0.15%) and those containing spurious lidar artifacts, identified by implausible reference heights (95th-percentile ≥ 97.13 m, mean > 62.0 m, or a 95th-to-mean difference ≥ 37 m; n = 775, 0.03%). Supplemental Figure S1 shows a random selection of reference tiles excluded due to these spurious lidar artifacts.

#### Evaluation Metrics

We computed mean absolute error (MAE), relative mean absolute error (RMAE), root mean square error (RMSE), bias, and the coefficient of determination (r^2^) using standard methods. Linear fits between predicted and reference heights were calculated using Reduced Major Axis (RMA) regression to account for measurement error in both variables. In addition to these standard metrics, we quantified the similarity between predicted and observed height distributions using the Jensen–Shannon Divergence (JSD; **equation 1**), a symmetric measure of distributional agreement:

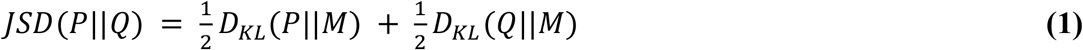

where

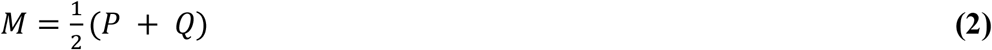

where *P* and *Q* are the normalized histograms of predicted and observed heights respectively, *M* is their average distribution (**equation 2**), and *D*_*KL*_ denotes the Kullback-Leibler divergence (**equation 3)**:

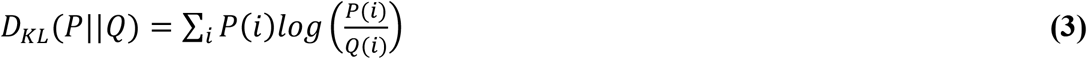

Height distributions were discretized into 60 bins spanning 0-60m (1m bin width), with a small constant (ε = 10^−10^) added to prevent numerical instability. JSD values are bounded between 0 and 1, with lower values indicating greater agreement between predicted and observed distributions. This metric provides insight into whether the model captures the full variability in canopy height beyond point- and tile-wise accuracy.

#### Post hoc error analysis

To understand where the model performs well and where it does not, we analyzed how tile-level prediction error varies with scene characteristics. We modeled log-transformed tile MAE as a function of three candidate predictors: observed canopy height (continuous), NLCD land-cover class, and EPA Level III ecoregion (categorical). We first compared univariate models to select the most informative canopy height metric (tile mean vs. 95th percentile). The selected height metric was then included as a fixed effect in linear mixed-effects models with random intercepts for (1) NLCD class, (2) ecoregion, or (3) both. We used variance partitioning^29^ to quantify the relative contribution each predictor to variation in prediction error across tiles. The purpose of this analysis is not causal attribution, but rather to provide practical context for users interpreting the patterns of model error visible in Figures 5 and 6 — specifically, whether apparent geographic variation in accuracy reflects systematic regional biases or is primarily a consequence of the well-documented relationship between canopy height and prediction error in CNN-based models.

### CONUS-wide Dataset Generation

Using the most recent NAIP imagery available, we produced a canopy height dataset across CONUS at 0.6-meter resolution (**Fig. 3)**. We used NAIP Digital Ortho Quarter Quads (DOQQs) as the primary unit of inference. DOQQs were retrieved using Google Earth Engine^30^ and split into 432×432 pixel tiles with 20% overlap. In all, we processed 213,567 DOQQs, covering approximately 7.7 million km^2^. The resulting CHMs were saved as Cloud-Optimized GeoTIFFs (COGs) with DEFLATE compression and internal overviews. The complete dataset with accompanying metadata totals approximately 24.82 TB.

**Figure 3:**
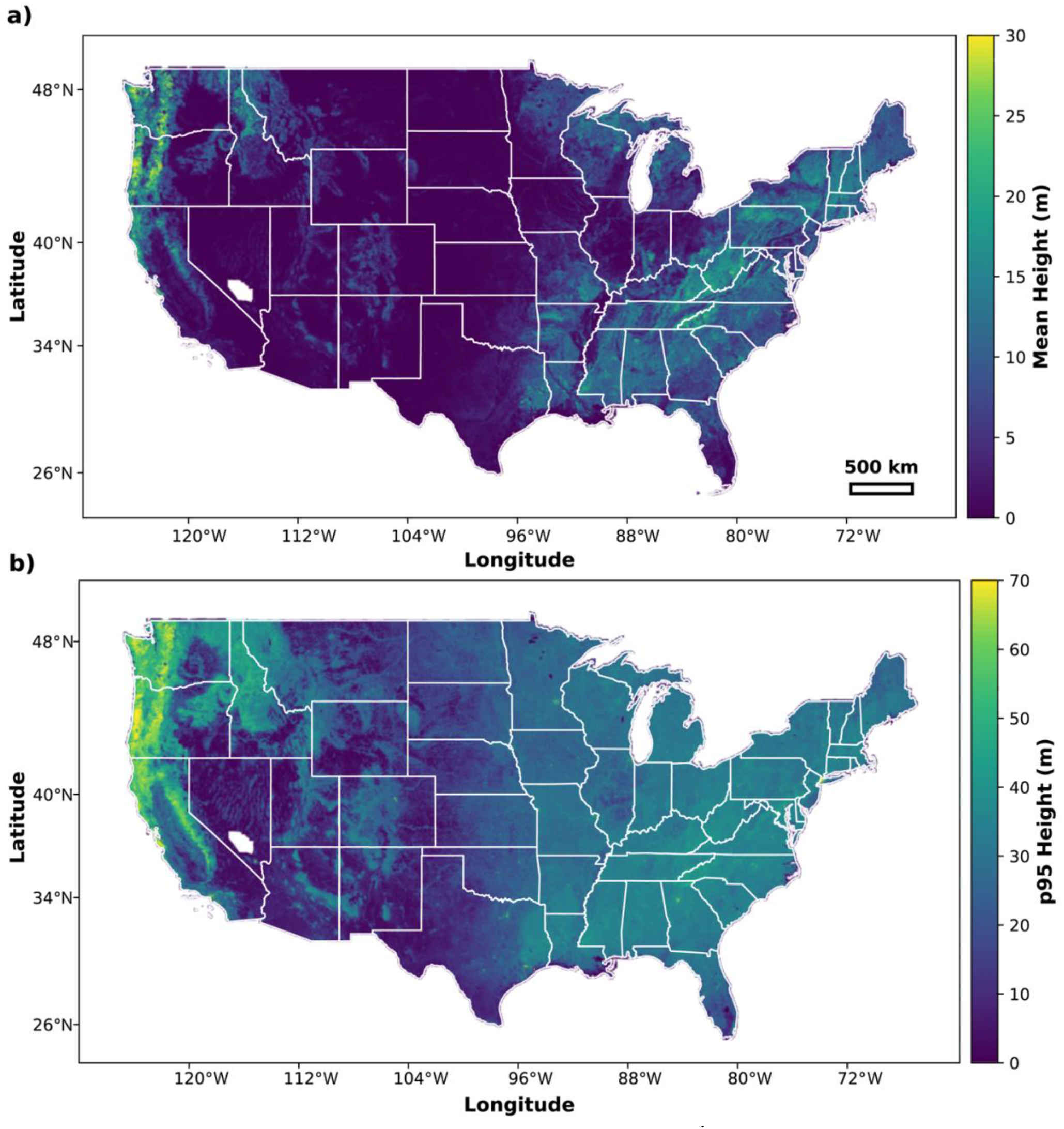
Modeled canopy height. Maps display the (a) mean and (b) the 95^th^ percentile (p95) canopy height derived from the NAIP-CHM. The void in coverage in southern Nevada is the Nevada Test and Training Range, where NAIP imagery is restricted.

### Data Records

The NAIP-CHM dataset, model, and associated code are distributed across several platforms to ensure accessibility and usability for different applications.

#### 1. CONUS Canopy Height Dataset

The NAIP-CHM 0.6-meter dataset is publicly available through multiple access points. For bulk download, COGs and associated index files are available via HTTP from the Rangeland Analysis Platform server (*http://rangeland.ntsg.umt.edu/data/naip-chm/*). Access through Google Earth Engine (ImageCollection ID: *projects/naip-chm/assets/conus-structure-model*) facilitates interactive analysis, particularly for non-specialists. For programmatic cloud-based workflows, COGs are also available in a public Google Cloud Storage bucket (*gs://naip-chm-assets/*) with “requester pays” enabled. In addition to the data, we provide a simple, interactive web application for data visualization and exploration, found at *https://naip-chm.projects.earthengine.app/view/naip-chm-a-conus-structure-model*.

#### 2. Model and Code Archive

A version-of-record archive of the model code, model weights, validation data, and auxiliary data used to generate the results presented in this manuscript is deposited in the Zenodo repository^31^.

#### 3. Development Repository and Code

The full codebase for model training, inference, and data processing is available in GitHub (https://github.com/smorf-ntsg/naip-chm). This repository includes examples demonstrating how to apply the model to future or past NAIP imagery using a Google Colab notebook and the Earth Engine API.

### Technical Validation

Performance was evaluated against the test partition of the published lidar-NAIP dataset (n = 2,273,381 tiles), which was held out entirely during model training and development. Pixel-level mean absolute error (MAE) was 0.89 m, root mean square error (RMSE) was 2.28 m, and bias was -0.02 m (**Table 2**). The r^2^ was 0.87, and the Jensen-Shannon Divergence (JSD) was 0.011, indicating close correspondence between predicted and observed height distributions. **Figure 4** illustrates pixel-level prediction performance across three representative land cover types, selected at each class’s median test-set performance. **Table 2** reports metrics stratified by NLCD land cover class. Within forested areas, the model achieved a pooled r^2^ of 0.82, MAE of 2.28 m, RMSE of 3.82 m, and bias of 0.10 m. Additional performance metrics by cover class are provided in ***Supplement A***.

**Table 2:** Test-set accuracy by land cover class.

| Land Cover | n tiles | Reference<br>Mean Height<br>(m) | p95<br>Reference<br>Mean Height<br>(m) | MAE<br>(m) | rMAE<br>(pct) | RMSE<br>(m) | Bias<br>(m) | r<sup>2</sup> | JSD |
| --- | --- | --- | --- | --- | --- | --- | --- | --- | --- |
| Water | 17,919 | 2.741 | 9.949 | 0.803 | 30.3 | 2.376 | -0.045 | 0.863 | 0.0107 |
| Developed | 203,912 | 4.096 | 14.23 | 1.171 | 28.6 | 2.487 | -0.073 | 0.852 | 0.0086 |
| Barren | 39,077 | 0.668 | 2.839 | 0.333 | 49.8 | 1.513 | -0.091 | 0.691 | 0.0085 |
| Forest | 263,644 | 9.741 | 19.685 | 2.282 | 23.4 | 3.820 | 0.096 | 0.816 | 0.0236 |
| Shrubland | 531,174 | 1.773 | 5.91 | 0.600 | 33.9 | 1.695 | -0.064 | 0.832 | 0.0089 |
| Grassland | 476,733 | 1.492 | 4.831 | 0.510 | 34.2 | 1.822 | -0.051 | 0.832 | 0.0077 |
| Pasture/Hay | 330,476 | 2.508 | 11.07 | 0.687 | 27.4 | 1.914 | -0.011 | 0.886 | 0.0055 |
| Cultivated Crops | 237,606 | 1.110 | 5.255 | 0.353 | 31.9 | 1.341 | -0.015 | 0.879 | 0.0100 |
| Wetlands | 172,840 | 6.656 | 15.012 | 1.651 | 24.8 | 3.189 | 0.059 | 0.85 | 0.0188 |
| All Classes | 2,273,381 | 3.240 | 9.368 | 0.893 | 27.5 | 2.277 | -0.022 | 0.871 | 0.0107 |

**Note:** Statistics computed on a pixel-wise basis, except for JSD which was calculated per-tile. The original CHM reference data were at 1.0 m resolution but were resampled to ~0.6 m for modeling. See Figure S2 for examples of the land cover classes used in this model.

**Figure 4.**
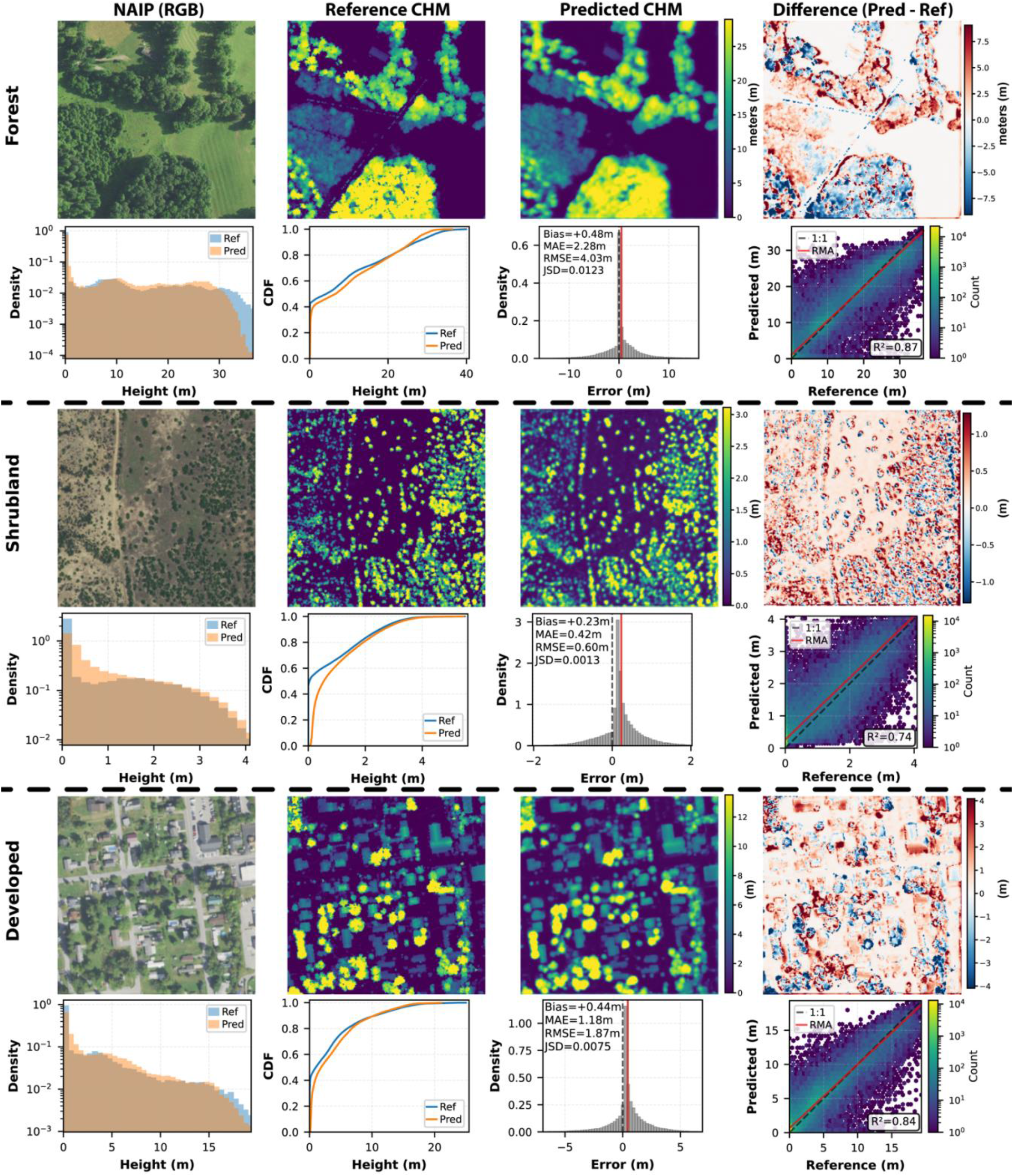
Pixel-level canopy height prediction performance across three representative land cover types (256 m x 256 m). For each class, the top row shows NAIP RGB imagery, lidar-derived reference CHM from our test dataset, model-predicted CHM, and the signed difference map. The bottom row presents pixel-level height distributions, cumulative distribution functions (CDF), error distributions with summary statistics, and predicted vs. reference scatter plots with a RMA regression fit line (red).

Aggregating at the tile level (256 × 256-meter patch size; n = 2,273,381), predicted mean and 95th percentile heights showed strong agreement with reference data (r^2^ = 0.98 and 0.97, respectively), with the RMA regression lines closely tracking the 1:1 identity line (**Fig. 5a, b**). The model exhibits a compression effect consistent with similar deep learning methods<u>^8–10^</u>, underestimating the tallest canopies by approximately 10%. Performance may be reduced in forests with canopies exceeding 70 m, such as those in the Pacific Northwest, as these areas were generally absent from the training data. Users interested in deriving biomass estimates from these canopy height predictions should note that this compression effect may propagate through nonlinear allometric relationships between height and biomass^32^, and additional calibration steps may be warranted^6,8^.

**Figure 5:**
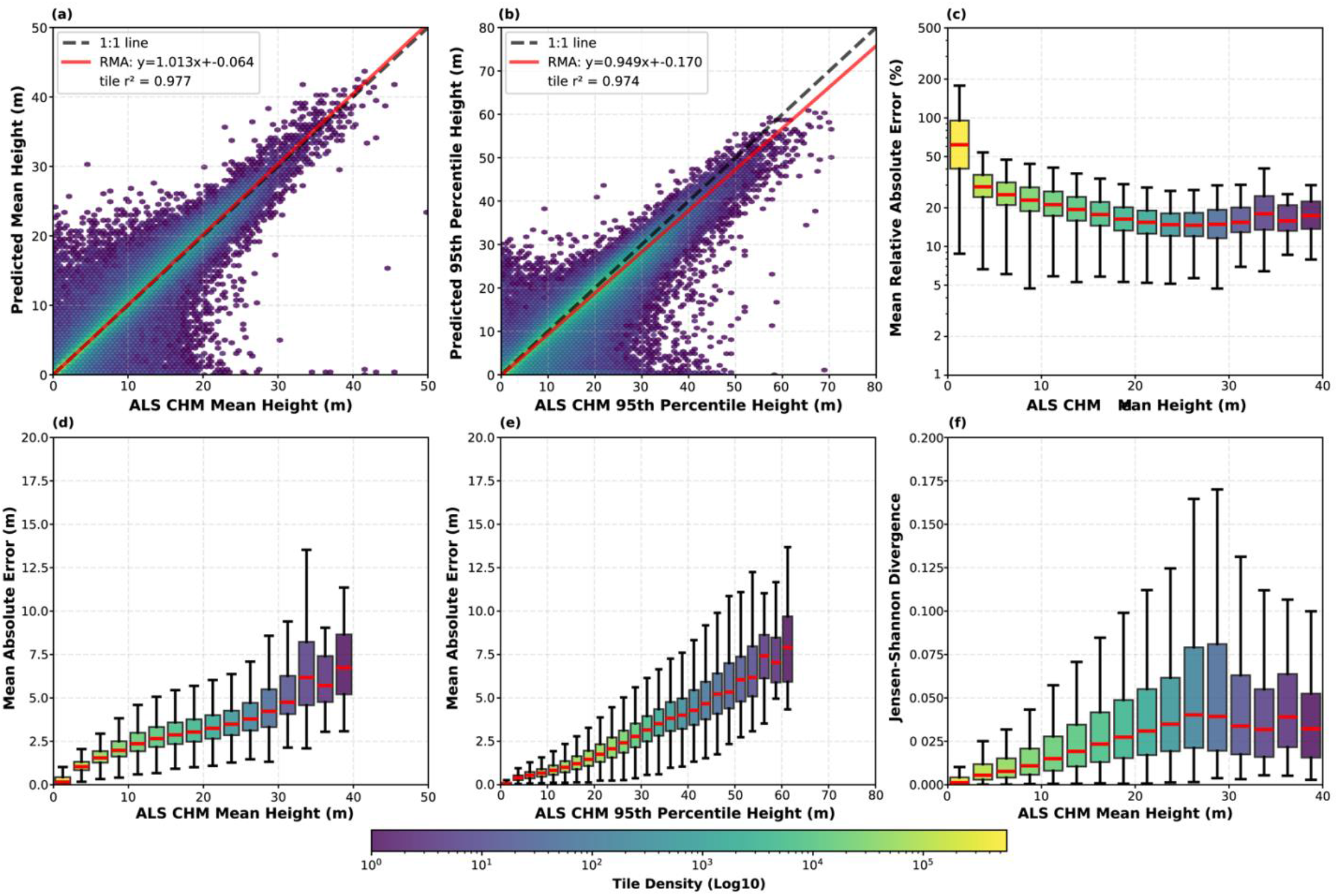
Tile-level validation of the NAIP derived Canopy Height Model (CHM) against the lidar derived CHM reference data. Panels (a) and (b) display density scatter plots for mean height and 95th percentile height, respectively, including the RMA fit (red solid line) and 1:1 identity (black dashed line). Panels (c–f) present box plots of error metrics binned by reference height: (c) Mean Relative Absolute Error in log-scale, (d) Mean Absolute Error (MAE) versus mean height, (e) MAE versus 95th percentile height, and (f) Jensen-Shannon Divergence. Red lines within the box plots indicate the median value.

Absolute error increased with canopy height, whereas relative error decreased (**Fig. 5c–e**). Analysis of the Jensen-Shannon Divergence shows that mean JSD increases only modestly with canopy height, confirming that the model accurately reproduces the reference height distributions within tiles across a range of canopy structures (**Fig. 5f**).

To characterize spatial variation in model performance, we decomposed tile-level prediction error (log MAE) using linear mixed-effects models. The 95th percentile of observed canopy height explained 59.3% of the variance in prediction error (marginal r^2^), reflecting the systematic underestimation of the tallest canopies common to deep-learning height models. Land cover and ecoregion together accounted for only 11.8% of the variance (conditional r^2^ = 0.71), with the remaining ∼29% likely reflecting tile-level factors such as temporal mismatch between NAIP and lidar acquisitions (e.g., leaf-on vs leaf-off lidar acquisitions) or variability in reference data quality. Because ecoregion and land cover influence canopy height, their total effect on prediction error is larger than 11.8% (∼47% via path analysis). However, this indirect path reflects the physical reality that taller canopies are harder to predict from optical imagery regardless of location — a compression artifact common to CNN-based height models, not a geographic bias. The direct effect of geographic context (<12%) indicates the model does not exhibit substantial systematic regional errors beyond what canopy height alone predicts, supporting generalizability across CONUS.

The geographic distribution of model performance reflects this error analysis (**Fig. 6**). Higher absolute errors (MAE) are concentrated in regions with tall canopies, such as the Pacific Northwest, northern Rocky Mountains, and Appalachians (**Fig. 6a**). Regional bias remains near zero, though a subtle pattern of positive bias in eastern forests and negative bias in western forests is apparent (**Fig. 6b**). The positive bias in eastern forests may partly reflect a phenological mismatch: lidar acquisitions in the eastern U.S. are more frequently collected during leaf-off conditions, whereas NAIP imagery is acquired during the growing season, leading the model to predict leaf-on canopy heights that exceed the leaf-off reference. r^2^ values are high (>0.8) across most ecoregions, and JSD remains low (<0.05), confirming consistent performance (**Fig. 6c, d**).

**Figure 6:**
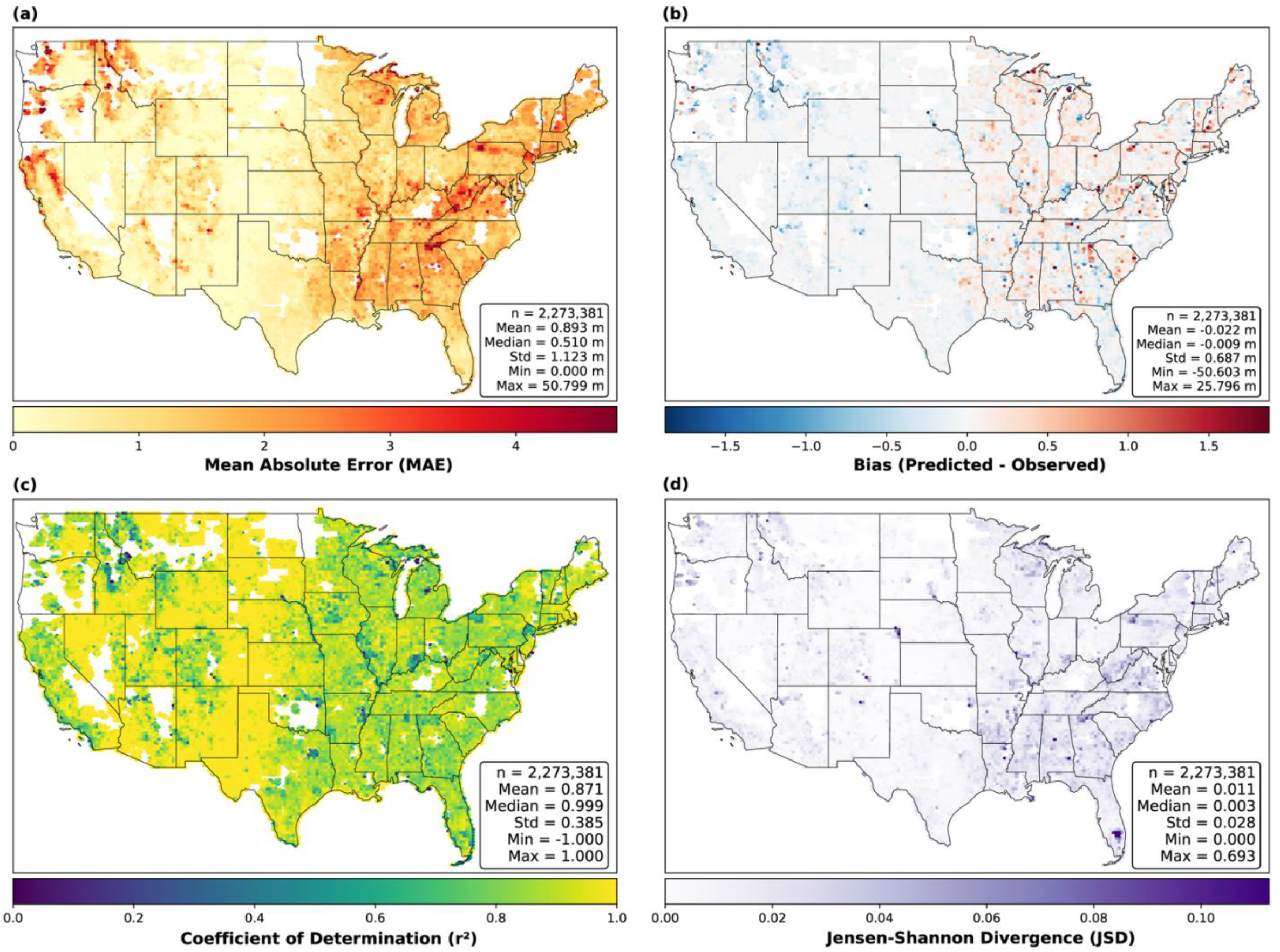
Geographic distribution of validation metrics across the contiguous United States. Panels display tile-level (a) Mean Absolute Error (MAE), (b) Bias, (c) Coefficient of Determination (r^2^), and (d) Jensen-Shannon Divergence (JSD). Inset boxes provide global summary statistics for each metric.

The difference between full-dataset and forest-only metrics illustrates how evaluation data distributions shape reported accuracies. Because prediction error scales with canopy height, CONUS-representative evaluations yield lower overall error than forest-only evaluations, which are restricted to tall-stature systems where absolute error is greatest. Readers comparing our reported metrics against other studies should account for differences in the height distributions of the underlying evaluation data.

To further characterize model generalization, we conducted two supplemental analyses evaluating performance across independent temporal and spatial holdouts. To test sensitivity to specific training acquisitions, we compared baseline prediction error (training-era NAIP vs. original lidar) against out-of-sample error (new NAIP vs. new lidar) at 228,123 matched locations (mean temporal separation: 4 years; **Supplement B**). When reference phenology is held constant (Leaf-On pairs), performance degradation is minimal (mean MAE and RMSE increased 0.08 m and 0.14 m, respectively, and r^2^ decreased by 0.02), and error distributions are nearly identical across training, validation, and test partitions — indicating the model is not dependent on the specific acquisitions seen during training.

To test whether performance degrades in landscapes absent from training, ten independent 10×10 km holdout regions spanning diverse CONUS ecoregions (18 –186 km from the nearest training tile) were evaluated against independently generated lidar reference CHMs (**Supplement C**). Across all ten spatial holdout regions, the model achieved an aggregate MAE of 1.66 m and an r^2^ of 0.80. However, nine of the ten regions produced errors highly consistent with regional and global test metrics (aggregate MAE = 1.31 m, r^2^ = 0.85). The sole outlier (ny_southeast; MAE = 4.83 m) exhibited systematic positive bias driven by a leaf-off lidar reference evaluated against leaf-on NAIP predictions—a phenological mismatch in the reference data, not a spatial generalization failure.

As an additional independent comparison, NAIP-CHM predictions were compared against CHMs produced by the National Ecological Observatory Network (NEON) Airborne Observation Platform^33^ at seven sites (**Fig. 7**). For this comparison, NAIP-CHM predictions were resampled to the 1.0 m resolution of the NEON reference CHMs. These products were generated using a separate acquisition and processing pipeline not used in model development; this comparison therefore assesses inter-product agreement rather than accuracy against ground truth. Across six of the seven sites, MAE ranged from 0.58 to 3.24-meter and r^2^ from 0.63 to 0.92. The seventh site (MOAB) was excluded because the NEON reference CHM contains widespread height values exceeding 10 m in a semi-arid landscape dominated by low stature Pinyon trees and shrubland — physically implausible values attributable to terrain model artifacts over steep, variable topography in the NEON model. Notably, image-based deep learning approaches like NAIP-CHM may be less susceptible to this failure mode, as they bypass the ground-classification step that limits traditional lidar CHM workflows in complex terrain.

**Figure 7:**
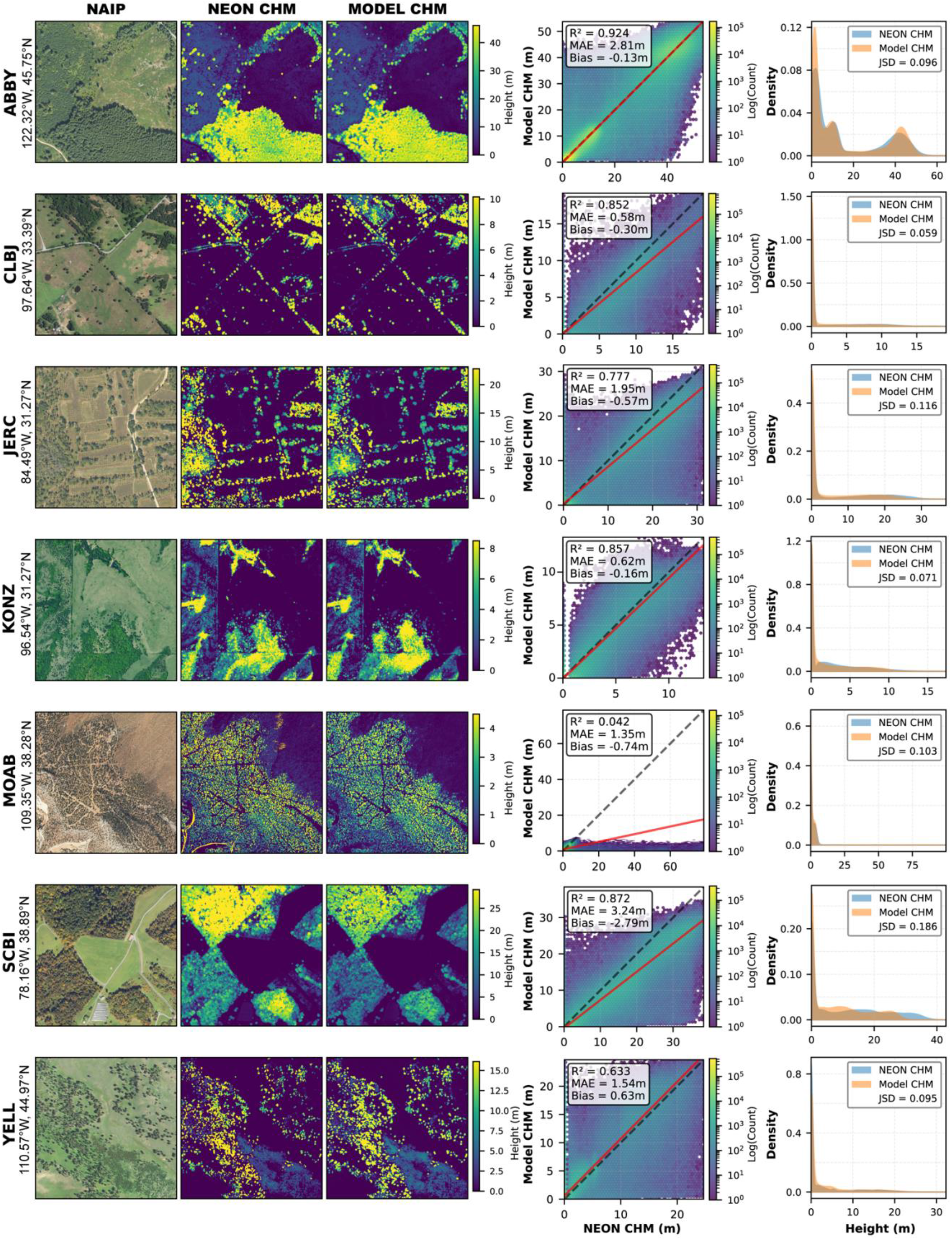
Comparison of NAIP-CHM predictions to National Ecological Observatory Network (NEON) canopy height models at seven sites, with predictions resampled to the 1.0 m NEON resolution. Scatter plots and histograms show r^2^, MAE, Bias, and Jensen-Shannon Divergence; dashed black lines indicate 1:1 and red lines show the RMA regression fit. Poor performance at MOAB reflects terrain artifacts in the NEON CHM. Visible data gaps at CLBJ and KONZ reflect NEON processing that zeros heights below 0.7–2.0 m. Sites: Abby Road (ABBY), Lyndon B. Johnson National Grassland (CLBJ), The Jones Center at Ichauway (JERC), Konza Prairie Biological Station (KONZ), Moab (MOAB), Smithsonian Conservation Biology Institute (SCBI), Yellowstone National Park (YELL).

Collectively, the temporal sensitivity, spatial holdout, and NEON analyses demonstrate that the model’s performance reflects learned physical relationships rather than spatial autocorrelation or acquisition-specific overfitting^27,34^. Although spatial cross-validation is frequently proposed to test for such overfitting, geographic blocking would artificially depress accuracy by withholding the continuous environmental covariates required by our FiLM conditioning layers^26,28^. Instead, our evaluation design relies on scale: with ∼3.4 × 10^12^ pixel-level regression targets and only 2.2 × 10^7^ trainable parameters, the model operates in a severely underdetermined regime (p/n ≈ 7 × 10^−6^) that severely limits finite-sample memorization^35,36^. Across independent temporal epochs, unseen geographic regions, and external processing pipelines, performance remains consistent with global metrics; where deviations do occur, they trace to canopy structure, phenological mismatch, or terrain-induced lidar artifacts rather than to spatial autocorrelation or acquisition-specific overfitting.

### Temporal applications

NAIP imagery is generally collected on a 2-3 year cycle. Although the NAIP-CHM dataset produced herein relied on the most recent NAIP imagery to produce a single snapshot of canopy height and structure, the model can be applied to both past and future NAIP 4-band imagery. This enables high resolution monitoring, capturing structural changes after disruptions or change events (**see Figure 8**), highlighting the model’s advantage for both broad-scale inventory *and* site-specific evaluation.

**Figure 8:**
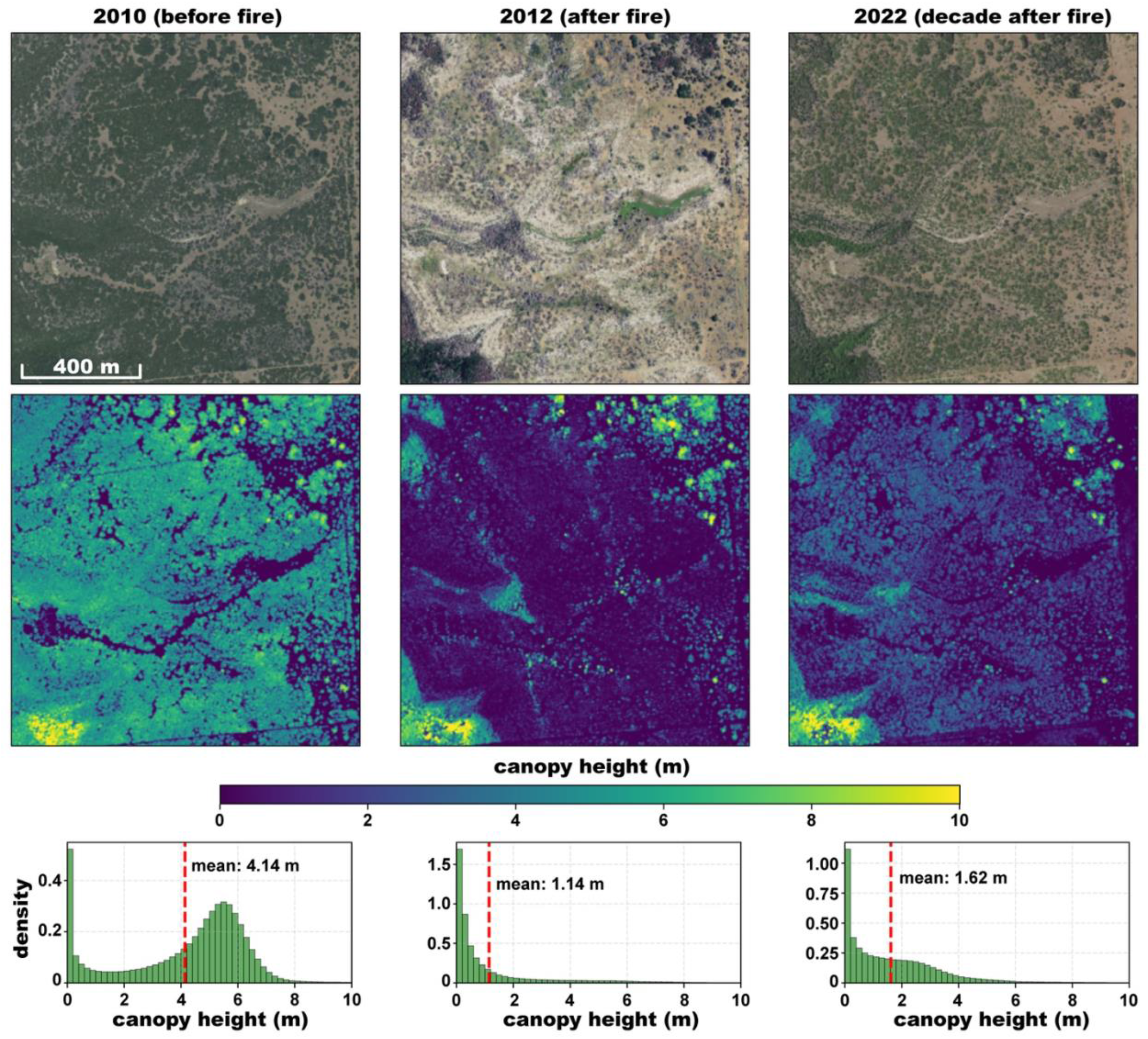
Chronosequence showing vegetation recovery following 2011 wildfire in central Texas (latitude N32.58, longitude W98.61). (a) Study area in 2010 (before fire), (b) 2012 (after 2011 fire), and (c) 2022 (∼ decade after fire). Middle row: CHMs showing the loss of canopy structure immediately following the fire event and subsequent regeneration. The density distributions show the shift from a closed woodland height structure (2010) to predominantly low vegetation (2012) and partial recovery (2022).

## Data Availability

The 0.6-meter resolution CONUS canopy height dataset is publicly available for bulk download via HTTP from the Rangeland Analysis Platform server (http://rangeland.ntsg.umt.edu/data/naip-chm/), as a Google Earth Engine asset (ImageCollection ID: projects/naip-chm/assets/conus-structure-model), and as Cloud-Optimized GeoTIFFs in Google Cloud Storage requestor-pays bucket (*gs://naip-chm-assets*).

## Code Availability

The source code, trained model weights, validation data, and auxiliary datasets required to reproduce the results are permanently archived in a Zenodo repository^31^. The active development repository, which contains scripts for model training, inference, and validation, is hosted on GitHub (https://github.com/smorf-ntsg/naip-chm) under an MIT License. To facilitate application of the model without local computational resources, we provide a Google Colab notebook (https://colab.research.google.com/github/smorf-ntsg/naip-chm/blob/main/notebooks/gee_inference_colab.ipynb) that allows users to run inference on past or future NAIP DOQQs.

## Acknowledgements

We thank Kris Mueller for assistance in figure preparation. This research used compute resources provided by (1) the SCINet project and/or the AI Center of Excellence of the USDA Agricultural Research Service (ARS project numbers 0201-88888-003-000D and 0201-88888-002-000D); (2) Google Earth Engine; (3) the Numerical Terradynamic Simulation Group at the University of Montana; and (4) the University of Montana’s Hellgate Research Cluster.

## Author contributions

S.L.M. led the conceptualization, methodology, software development, data curation, formal analysis, and visualization (web application), and wrote the original draft. B.W.A. contributed to conceptualization, software (model inference, code review), and validation. A.A.M. contributed to validation and software (code review). J.T.S. contributed to the formal analysis. D.E.N. provided supervision. All authors reviewed the model outputs, provided feedback during model development, and contributed to the review and editing of the manuscript.

