## Supplemental Materials for "A 0.6-meter resolution canopy height model for the contiguous United States"

**For the manuscript:**

---

**Overview of Supplementary Information:** This document provides supplementary figures, expanded error analyses, and independent validation assessments to support the data and methodology presented in the main manuscript. The material is organized into the following sections:

**Supplemental Figures (S1 & S2):** Provides visual context for the data processing and evaluation, including randomly selected examples of reference tiles excluded from the test dataset due to severe anomalous lidar noise (Figure S1), and representative 0.6-meter NAIP imagery and canopy height model (CHM) predictions for nine distinct National Land Cover Database (NLCD) classes (Figure S2).

**Supplement A: Model Performance Metrics by Land Cover Class**
Expands upon the summary metrics provided in the main text (Table 2). This section includes detailed, multi-panel visual breakdowns of model performance across nine aggregated NLCD land cover classes. It features pixel-level height and error distributions, tile-level regressions, error trends, and hexbin maps of spatial performance across the contiguous United States (CONUS) (Figures A1–A10).

**Supplement B: Temporal Acquisition Sensitivity Analysis**
Evaluates the model's robustness to temporal variation and its sensitivity to the specific image acquisitions used during training. By analyzing over 228,000 matched geographic locations with independent out-of-sample NAIP and lidar pairs, this section demonstrates that model accuracy is driven by learned canopy structure relationships rather than spatial autocorrelation or acquisition-specific overfitting (Figures B1–B4).

**Supplement C: Spatial Generalization**
Assesses the model’s ability to generalize to landscapes entirely absent from the training data. Performance is evaluated across ten geographically isolated 10 × 10 km holdout regions distributed across diverse CONUS ecoregions (ranging from arid shrubland to dense coastal conifer forests), demonstrating reliable generalization limited only by expected physical and phenological constraints (Tables C1–C2, Figures C1–C3).

#### Supplemental Figures:

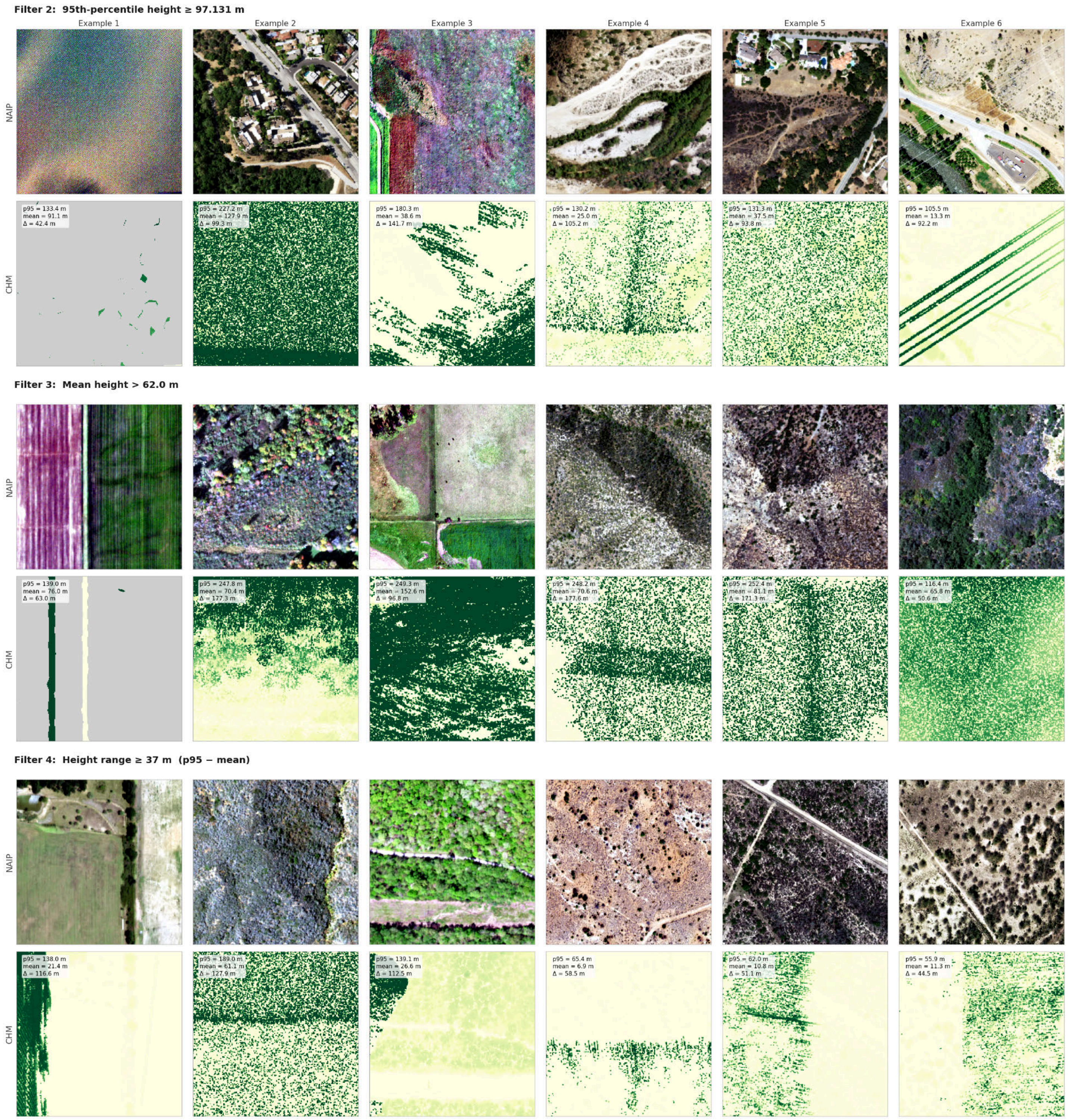


**Figure S1:** Randomly selected examples of reference tile pairs excluded from the test dataset due to anomalous reference heights. Each pair shows the NAIP imagery (top) alongside the reference canopy height model (CHM) (bottom). This heuristic removed 775 tile pairs (0.03% of the test set) containing severe elevation noise in the lidar reference data that would otherwise skew validation metrics. Text annotations on the reference CHMs denote the 95th-percentile reference height (p95), the mean reference height (mean), and their difference (delta).

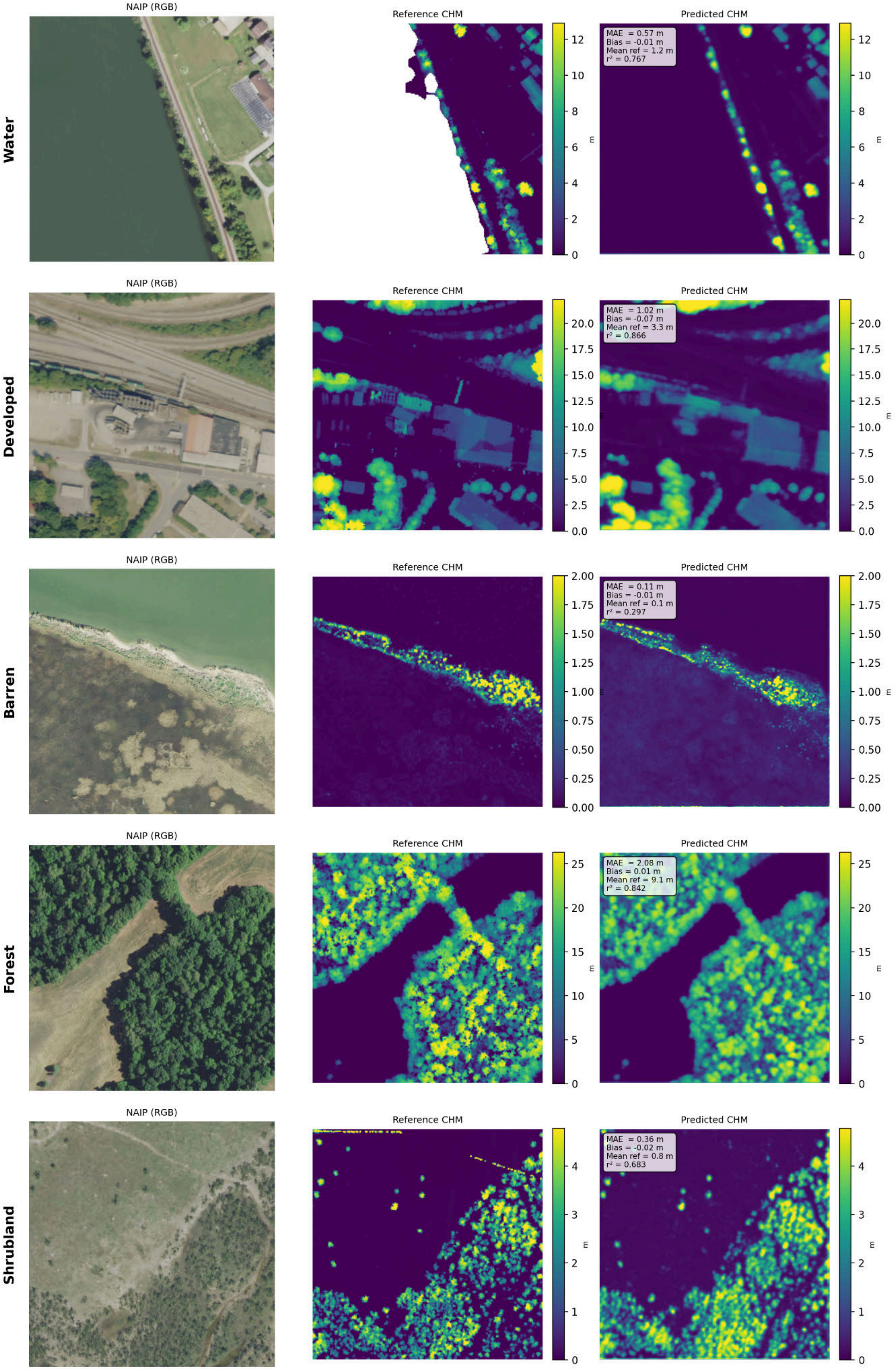
Water
Developed
Barren
Forest
Shrubland
NAIP (RGB)
Reference CHM
Predicted CHM
MAE = 0.57 m
Bias = -0.01 m
Mean ref = 1.2 m
r² = 0.767
MAE = 1.02 m
Bias = -0.07 m
Mean ref = 3.3 m
r² = 0.866
MAE = 0.11 m
Bias = -0.01 m
Mean ref = 0.1 m
r² = 0.297
MAE = 2.08 m
Bias = 0.01 m
Mean ref = 9.1 m
r² = 0.842
MAE = 0.36 m
Bias = -0.02 m
Mean ref = 0.8 m
r² = 0.683
m

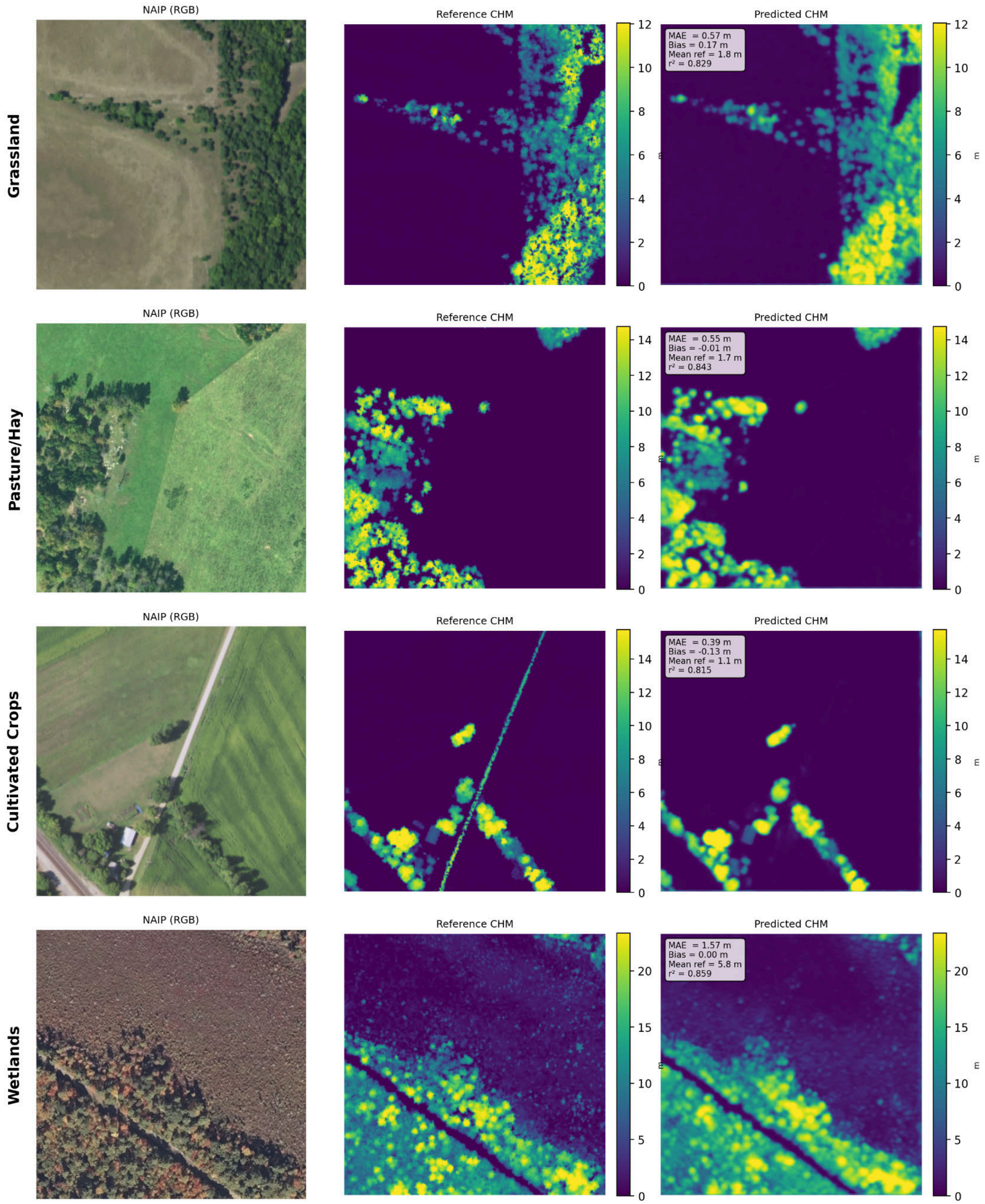


**Figure S2:** Representative NAIP tiles for each NLCD land cover class based on median performance statistics:(left) NAIP true-color imagery, (center) reference canopy height model derived from airborne lidar, and (right) predicted canopy height model from the UNetFiLM model.

#### Supplement A: Model Performance Metrics by Land Cover Class

##### Overview

As noted in the main text, overall and class-stratified summary statistics are provided in Table 2. This supplement expands upon those summary metrics by providing detailed, multi-panel visual breakdowns of model performance for each of the nine aggregated land cover classes. Following the classification scheme established in the lidar-NAIP dataset, National Land Cover Database (NLCD) codes are aggregated into nine distinct groups: Water (NLCD codes 11, 12), Developed (21–24), Barren (31), Forest (41–43), Shrubland (51, 52), Grassland (71–74), Pasture/Hay (81), Cultivated Crops (82), and Wetlands (90, 95).

The tile-level quality filters detailed in the main text—specifically, the minimum valid-pixel fraction, reference 95th percentile (P95) and mean height caps, and the P95–mean range cap—are applied uniformly across all classes. Pixel-level metrics are calculated by pooling all valid pixels within a given cover class, whereas per-tile metrics are derived from aggregated statistics at the tile level (256 × 256 m).

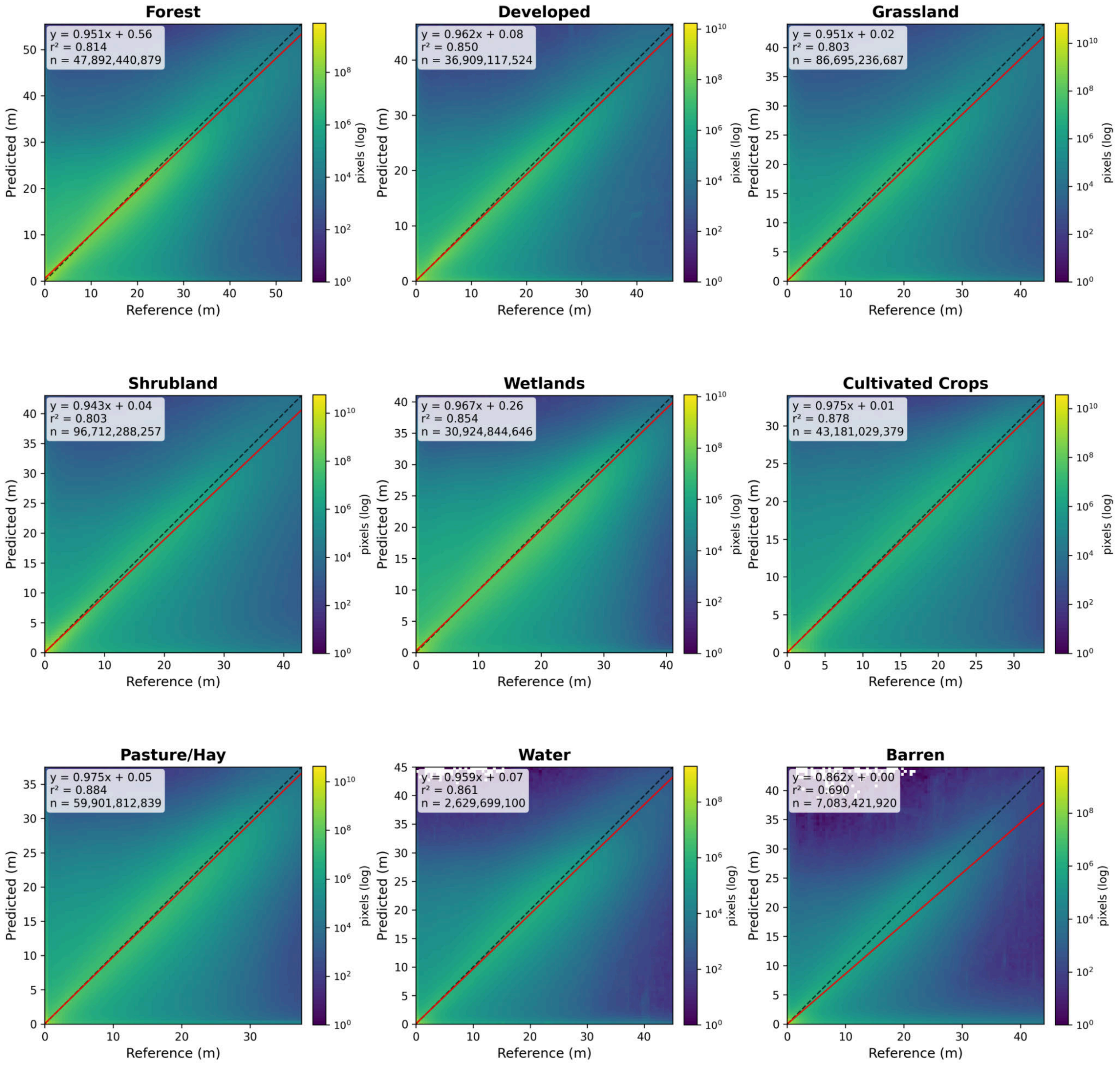


**Figure A1. Pixel-level canopy height regression by cover class.** Two-dimensional histograms illustrate the relationship between predicted and reference (ALS) heights for each of the nine land cover classes. Logarithmic color scales are employed to emphasize both the bulk data distribution and the low-density tails. The solid red and dashed black lines represent the reduced-major-axis (RMA) regression and the 1:1 identity line, respectively. Insets within each panel detail the regression equation (calculated from accumulated pixel moments), the pixel-wise coefficient of determination (r2), and the total valid pixel count (n).

**Figures A2–A10. Comprehensive performance metrics per cover class.** For each of the nine cover-class groups, we provide a single-page, multi-panel summary figure (Figures A2–A10). These figures utilize the same structural components as those in the main text but are expanded to include pixel-level height and error distributions, as well as histograms of tile-level error (Mean Absolute Error [MAE] and Root Mean Square Error [RMSE]) and distribution divergence (Jensen–Shannon Divergence [JSD]).

##### Pixel- and Tile-Level Distributions (Panels a–f)

- **Panel (a) Pixel height distributions:** Overlaid histograms of reference (ALS) and predicted pixel heights. The y-axis is logarithmically scaled to highlight the tall-canopy tail, while the x-axis spans the 0 to 99.99th percentile range.
- **Panel (b) Pixel prediction error:** Histogram of signed residuals (predicted − reference). The x-axis is dynamically scaled per class to the symmetric 2.5th–97.5th percentile window of the residuals.
- **Panel (c) Pixel signed relative error:** Histogram of relative error, calculated as [(predicted − reference) / reference] × 100%. To ensure a well-defined denominator, this metric is restricted to pixels with a reference height ≥ 0.5 m. The distribution is bounded at −100% (as predictions are strictly non-negative) and plotted up to +100%.
- **Panel (d) Tile MAE and RMSE:** Overlaid histograms of tile-level Mean Absolute Error and Root Mean Square Error. The x-axis range extends to the 99th percentile of the combined metric values.
- **Panel (e) Tile P95 heights:** Overlaid histograms of reference and predicted tile-level 95th-percentile (P95) heights. The x-axis range extends to the 99th percentile of the combined metric values.
- **Panel (f) Jensen–Shannon divergence (JSD):** Histogram of the tile-level JSD between the reference and predicted pixel-height distributions within each tile. The x-axis range extends to the 99th percentile of the combined metric values.

##### Tile-Level Regressions and Error Trends (Panels g–l)

- **Panel (g) Tile-mean predicted vs. reference height:** Hexbin density plot comparing mean predicted and reference tile heights, featuring a dashed 1:1 line (black) and an RMA regression line (red).
- **Panel (h) Tile P95 predicted vs. reference height:** Hexbin density plot comparing predicted and reference 95th-percentile (P95) tile heights, featuring a dashed 1:1 line (black) and an RMA regression line (red).
- **Panel (i) Relative MAE vs. tile-mean reference height:** Boxplots of tile-level relative MAE (expressed as a percentage of the mean reference height), binned at 2.5 m intervals with Tukey whiskers.
- **Panel (j) Tile MAE vs. tile-mean reference height:** Boxplots of absolute MAE binned at 2.5 m intervals of mean reference height, with Tukey whiskers.
- **Panel (k) Tile MAE vs. tile P95 reference height:** Boxplots of absolute MAE binned at 2.5 m intervals of P95 reference height, with Tukey whiskers.
- **Panel (l) JSD vs. tile-mean reference height:** Boxplots of tile-level JSD binned at 2.5 m intervals of mean reference height, with Tukey whiskers.

##### Spatial Performance Across the Contiguous United States (CONUS) (Panels m–p)

Spatial distributions of model performance are visualized as hexbin maps on a Plate Carrée projection with overlaid state boundaries. Panels (m)–(o) represent metrics pooled across all valid pixels within a given spatial cell, whereas panel (p) represents the average of tile-level values within the cell.

- **Panel (m) Mean Absolute Error:** Pixel-pooled MAE.
- **Panel (n) Bias:** Pixel-pooled mean signed error (predicted − reference).
- **Panel (o) Coefficient of determination (r2):** Pixel-pooled r2, computed from summed pixel moments.
- **Panel (p) Jensen–Shannon divergence:** Per-cell mean of the tile-level JSD.

**Figure A2: Forest (n = 263,644)**

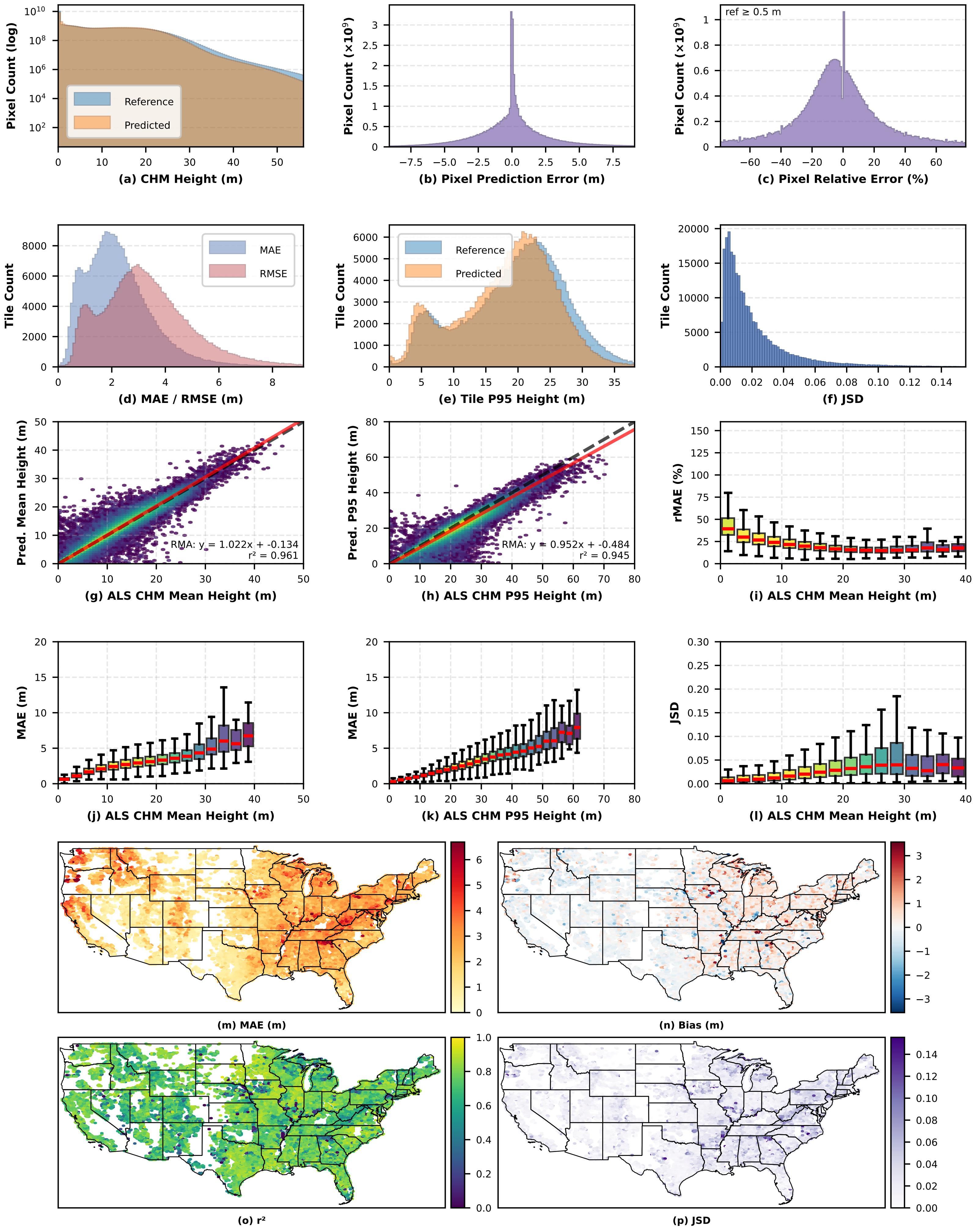

**Figure A3: Developed (n = 203,912)**

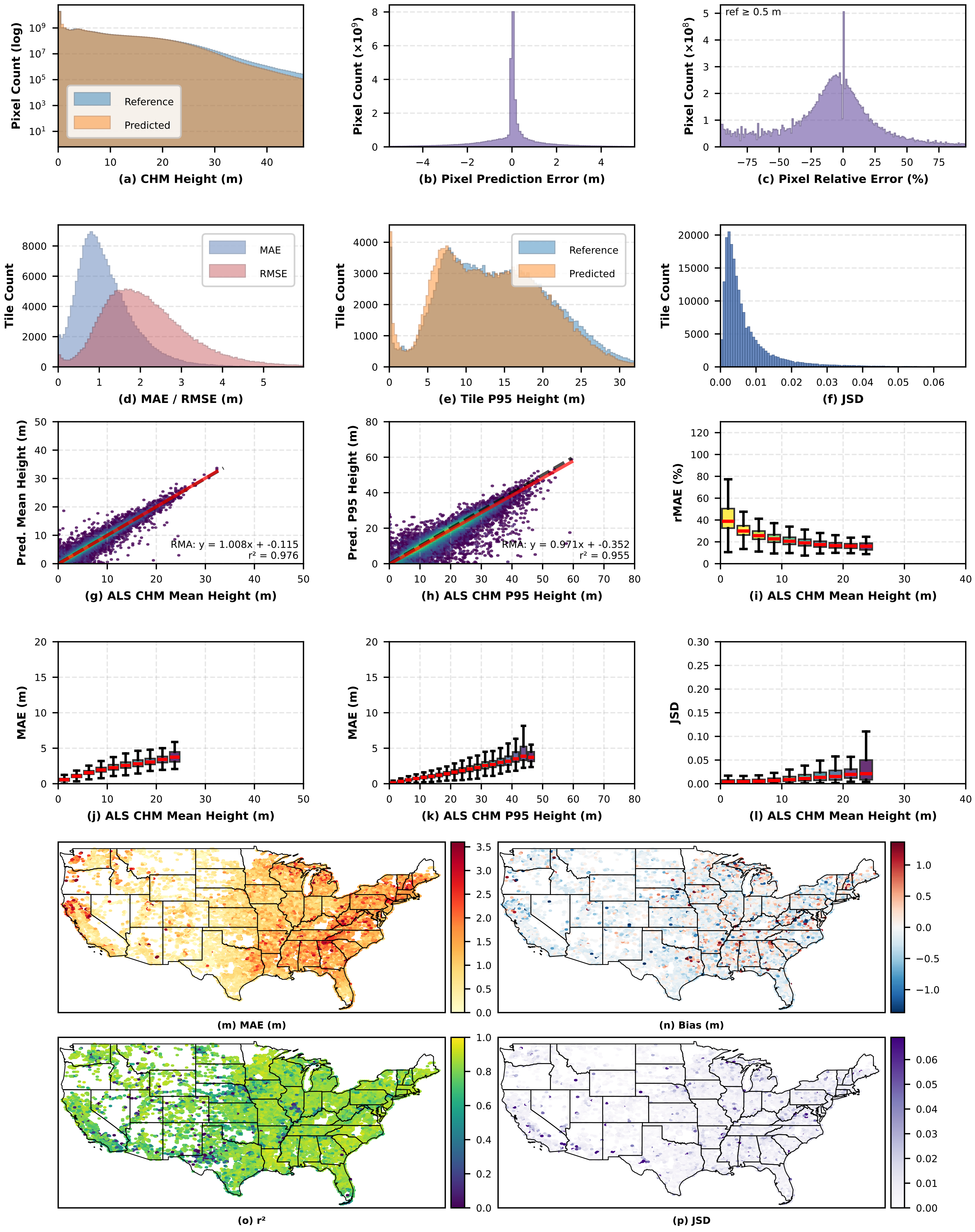

### Figure A4: Grassland (n = 476,733)

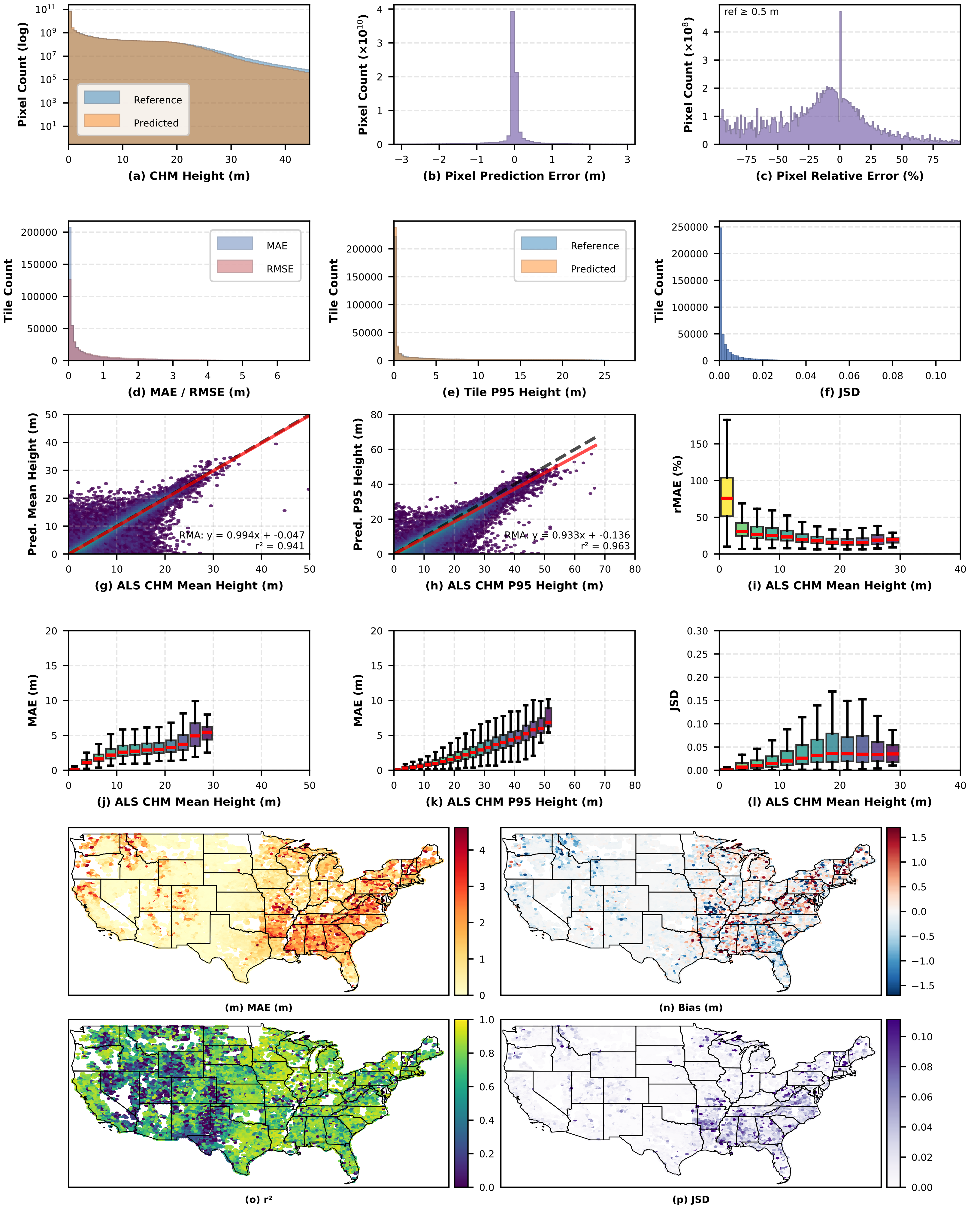

**Figure A5: Shrubland (n = 531,174)**

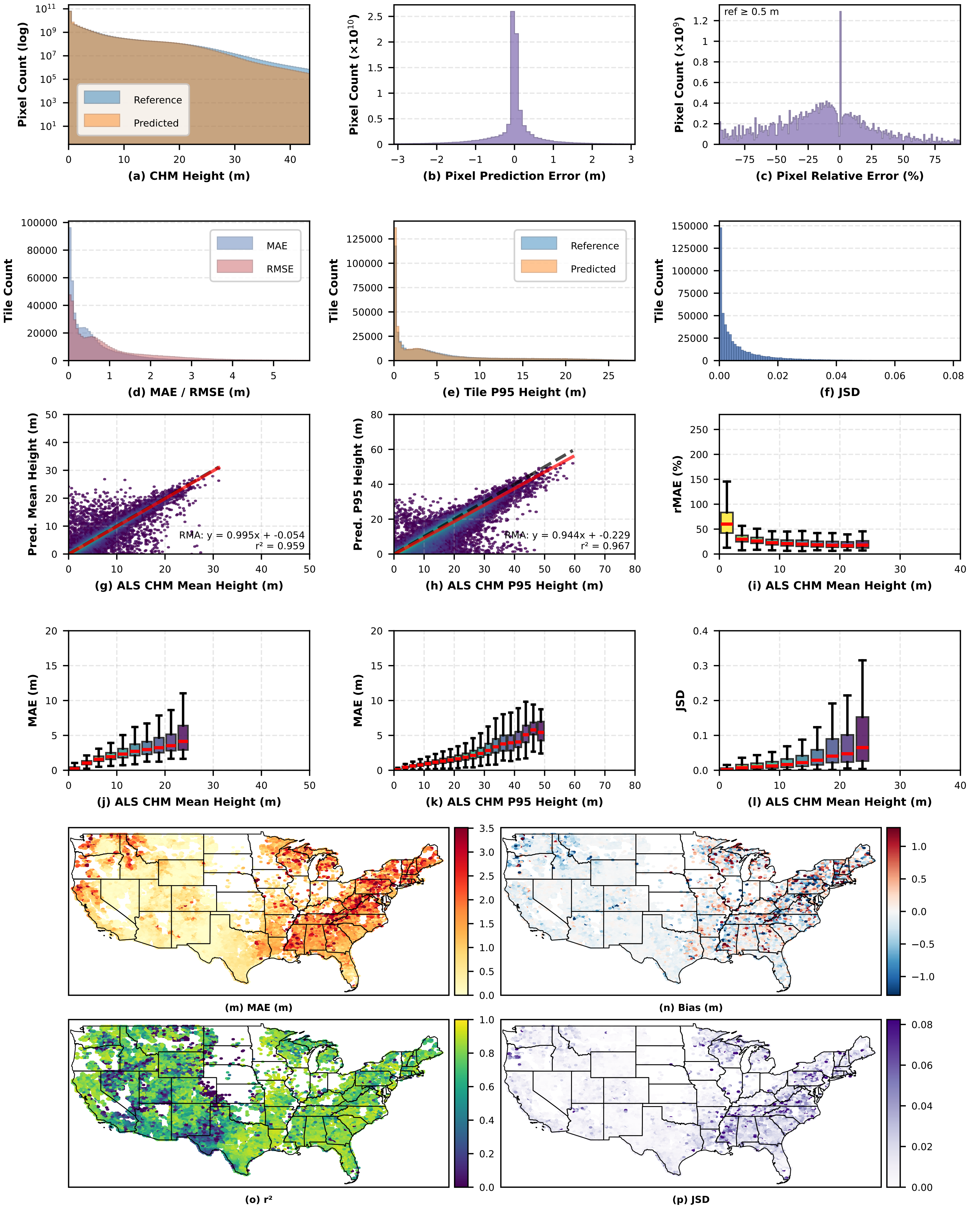

### Figure A6: Wetlands (n = 172,840)

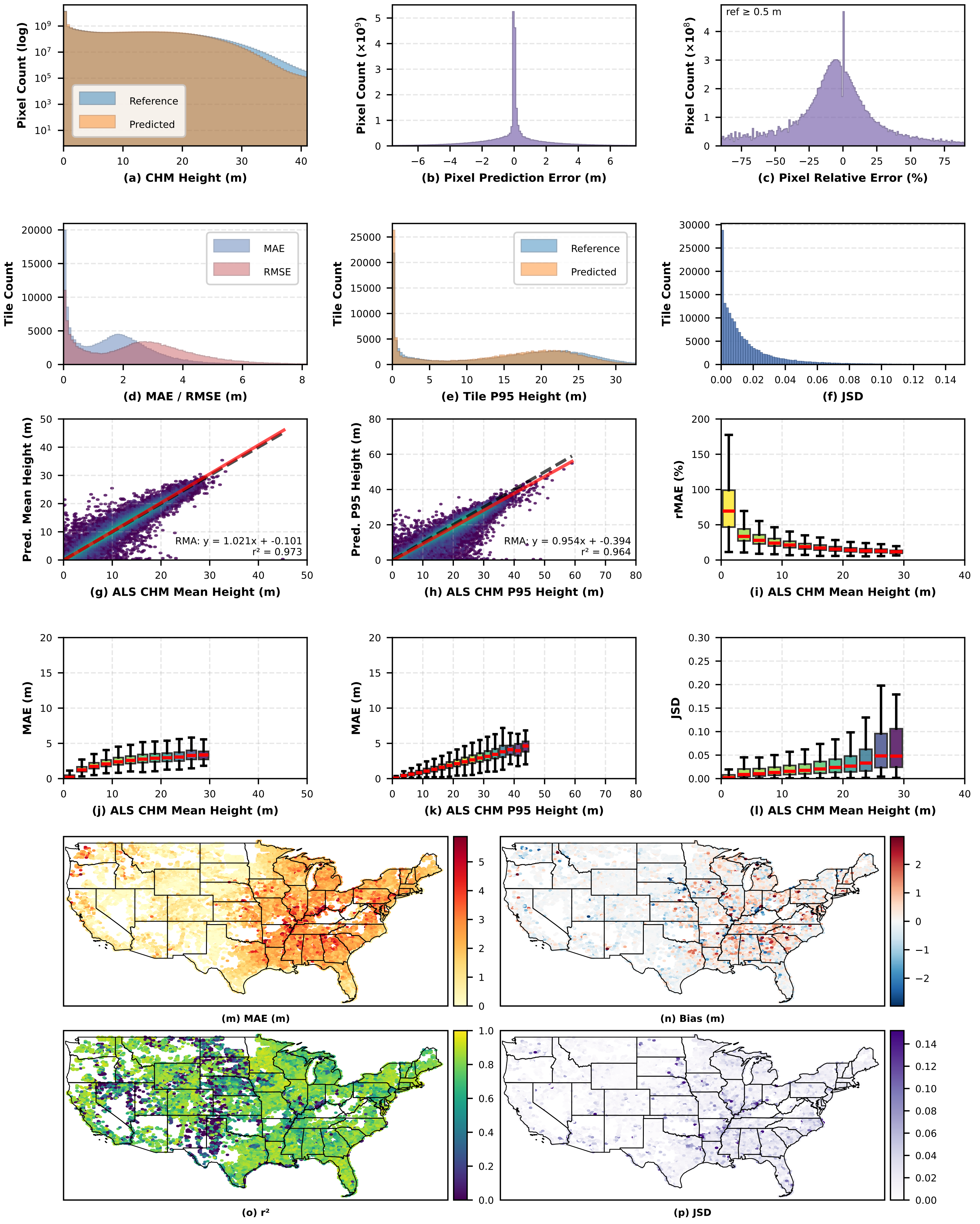

#### Figure A7: Cultivated Crops (n = 237,606)

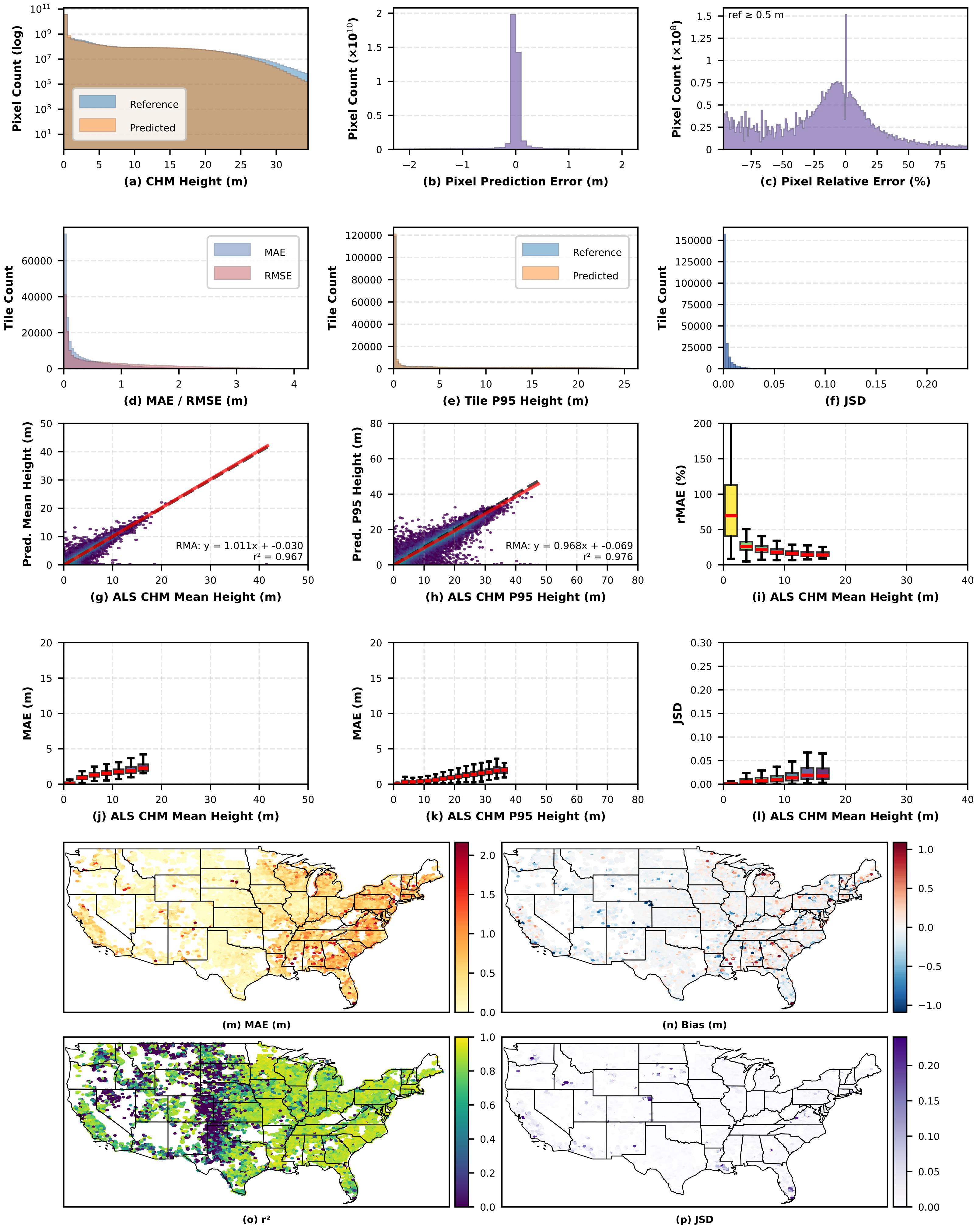

**Figure A8: Pasture/Hay (n = 330,476)**

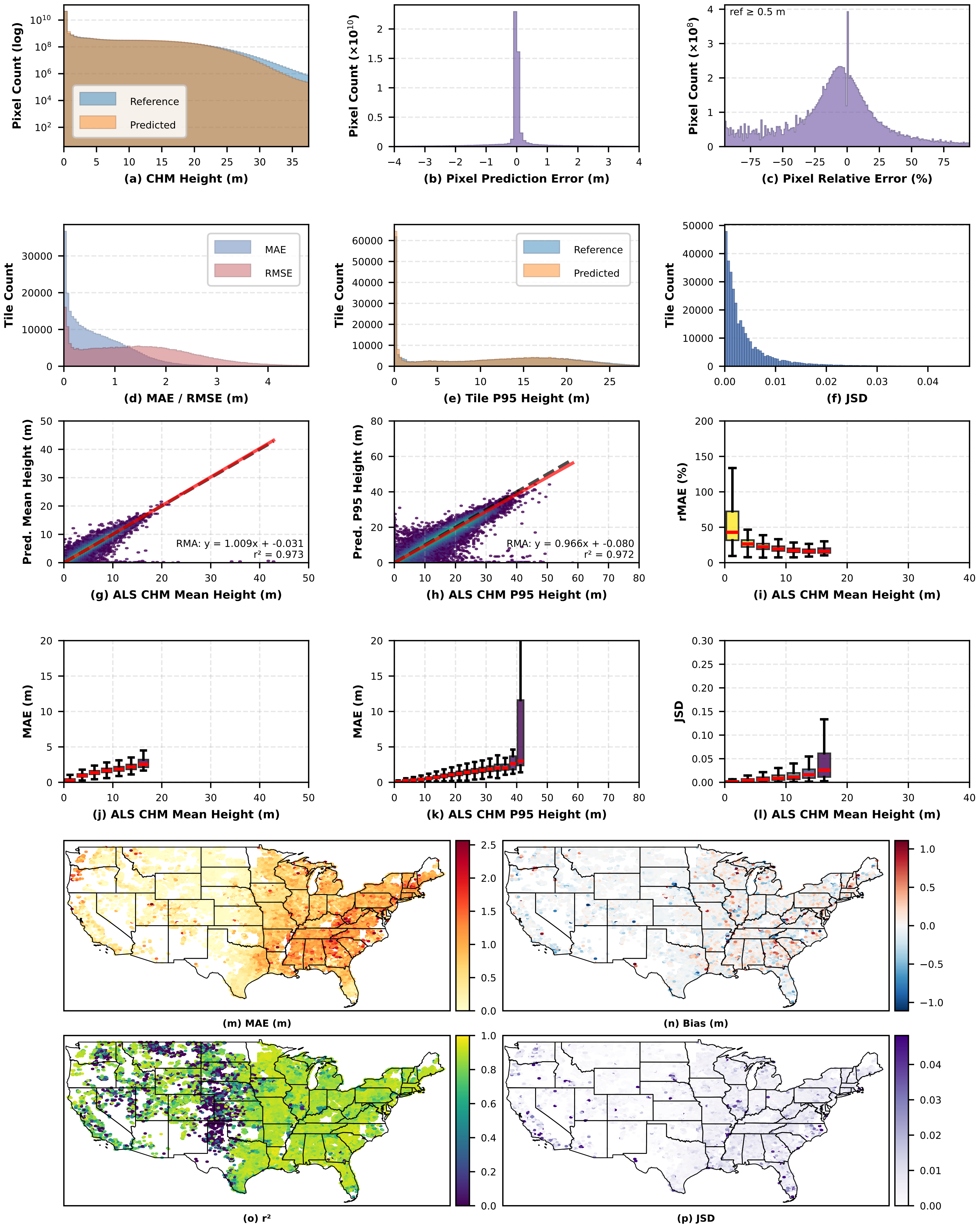

### Figure A9: Water (n = 17,919)

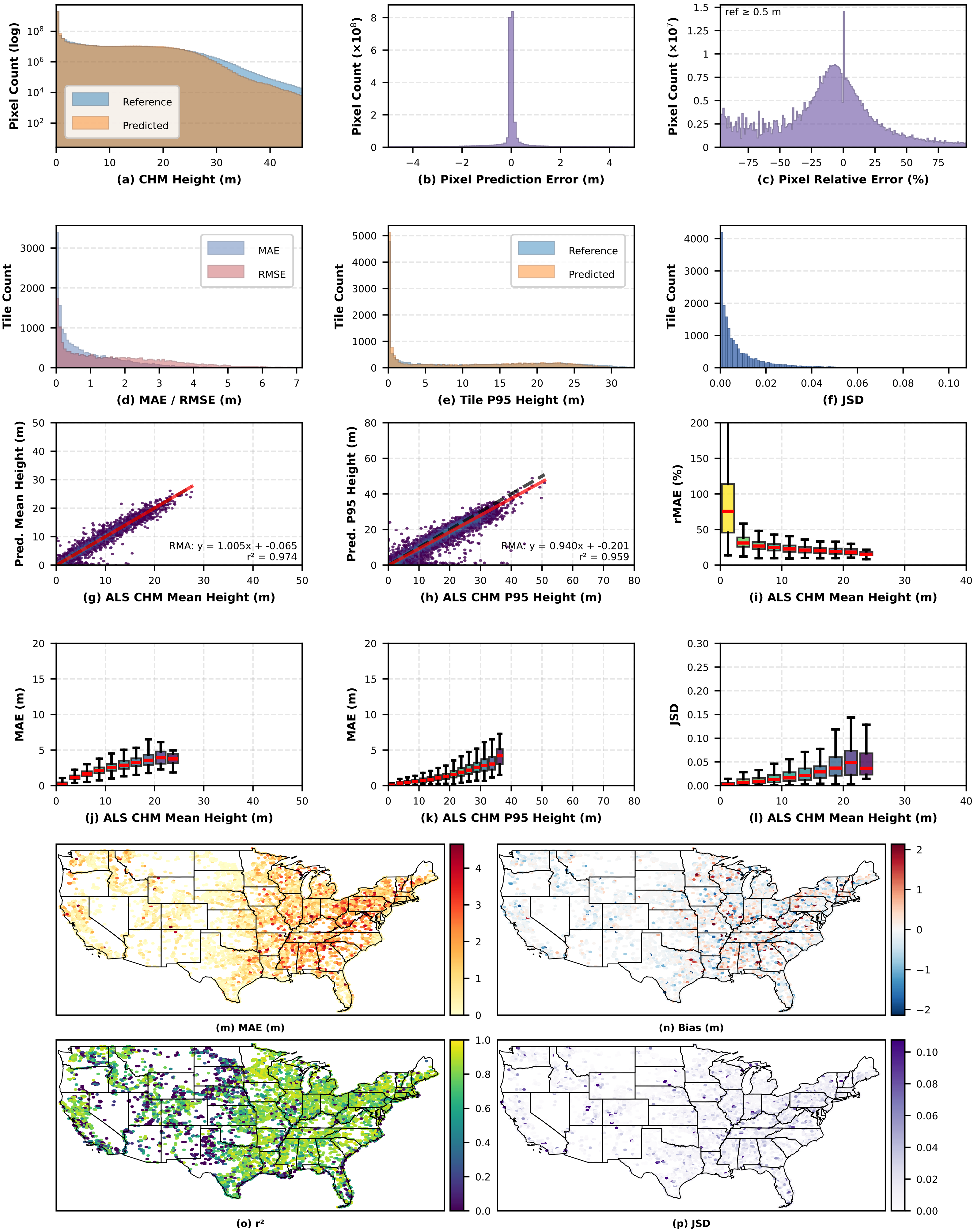

**Figure A10: Barren (n = 39,077)**

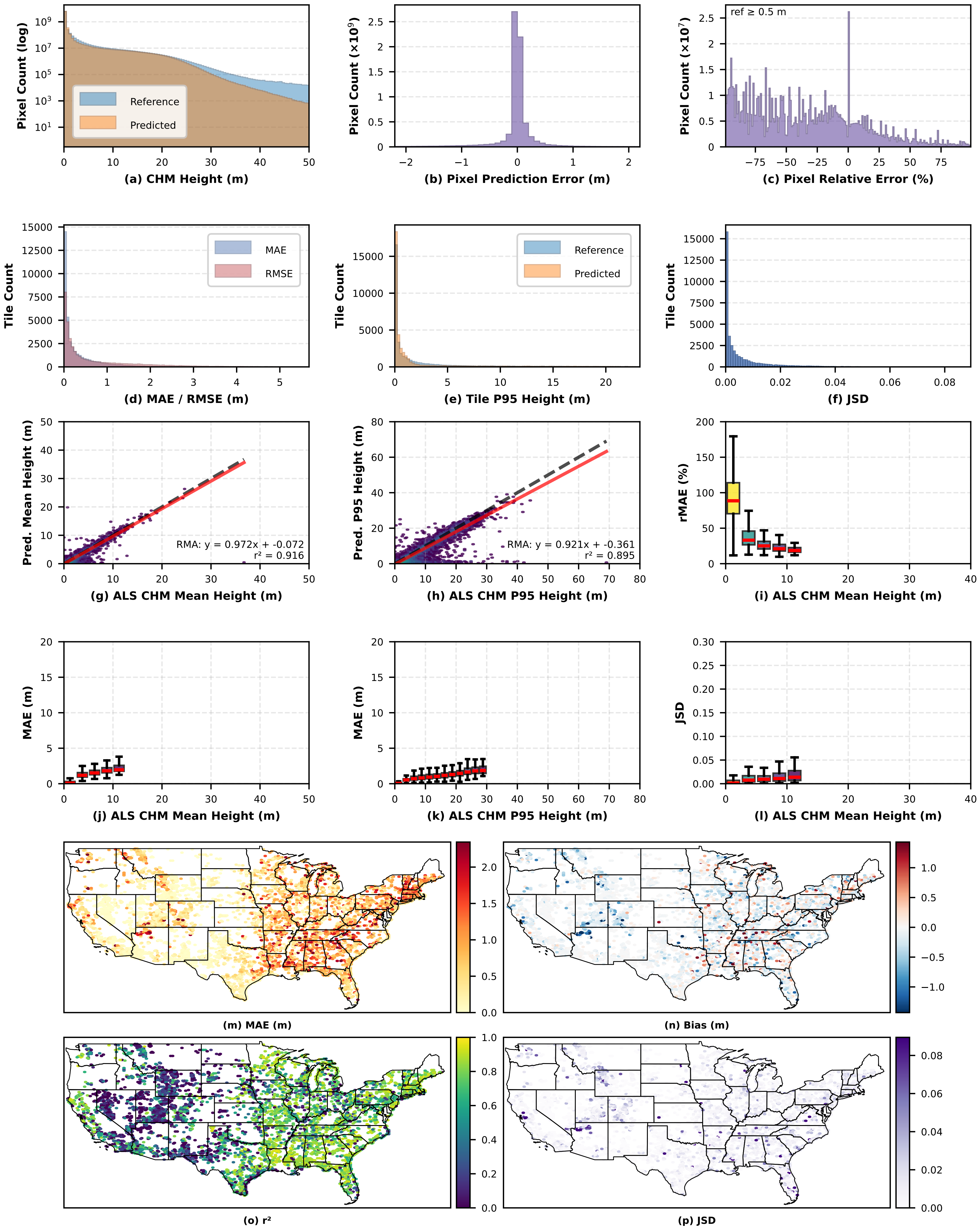

#### Supplement B: Temporal Acquisition Sensitivity Analysis

##### Overview and Methods

To evaluate model robustness to temporal variation in image acquisition, we compared baseline prediction error (training-era NAIP vs. original lidar) against fully out-of-sample error (new NAIP vs. new lidar) at 228,123 matched geographic locations (mean temporal separation: 4 years). This design tests whether model performance degrades when both the NAIP input and lidar reference are replaced with acquisitions never seen during model development.

Independent lidar point clouds for these locations were obtained from the USGS 3DEP Lidar Point Cloud archive[1,2], hosted publicly on Amazon Web Services in Entwine Point Tile (EPT) format (*s3://usgs-lidar-public*, AWS Registry of Open Data), with project footprints from the [hobuinc/usgs-lidar][3] (https://github.com/hobuinc/usgs-lidar) repository and acquisition dates from the USGS Work-unit Extent Spatial Metadata (WESM) layer of the 3DEP program.

New National Agriculture Imagery Program (NAIP)[4] acquisitions were identified and downloaded from the USDA Farm Production and Business Services NAIP Digital Ortho Quarter Quadrangle (DOQQ) collection via Google Earth Engine[5] (image collection `USDA/NAIP/DOQQ`). For each candidate location, we queried the collection for all NAIP acquisitions within ±3 years of the new lidar acquisition date and selected the temporally nearest scene. Four-band (R, G, B, NIR) image chips were then exported at native NAIP resolution (~0.6 m) matching the spatial extent of the corresponding lidar-derived CHM tile.

We identified 236,106 locations where independent lidar-derived CHMs and NAIP scenes were available from acquisitions not used in training. After applying quality filters consistent with our primary evaluation (minimum 25% valid pixels, 95th percentile height < 97.1 m, mean height < 62 m, height range < 37 m), 228,123 matched tile pairs remained.

Because lidar acquisitions span different seasons, phenological mismatch introduces a confounding factor unrelated to acquisition dependence: leaf-off lidar captures a reduced canopy surface relative to full-canopy predictions from leaf-on NAIP. We classified each lidar acquisition as leaf-on (mid-April to October) or leaf-off (November to March), producing four paired categories reflecting the phenological status of the original and new lidar references: On-On (n = 85,811), On-Off (n = 48,775), Off-On (n = 15,923), and Off-Off (n = 77,614). Their geographic distribution reflects regional differences in lidar acquisition timing, with western conifer regions predominantly On-On and eastern deciduous forests more frequently Off-Off or On-Off (**Fig. B1**). CHMs were processed from the raw lidar data using the same methods from the original dataset[6].

##### Results

*Performance by leaf-status category.* **Fig. B2** presents paired boxplots of bias, MAE, and JSD comparing training-year and non-training predictions within each category. The On-Off category exhibits the largest degradation, consistent with the model predicting full canopy from leaf-on NAIP while the leaf-off reference underrepresents canopy height. When leaf status is held constant, degradation is minimal: On-On pairs show a mean MAE increase of 0.07 m (from 1.22 to 1.29 m), mean RMSE increase of 0.14 m (from 2.11 to 2.25 m), $r^2$ decrease of 0.02 (from 0.84 to 0.82), and mean JSD increase of 0.004 (from 0.010 to 0.014). When pooling all categories and excluding only the severe On-Off phenological mismatch, overall performance degradation between training-year and non-training-year predictions remains similarly negligible (**Fig. B3**).

*Consistency across data partitions:* Within each leaf-status category, error distributions are nearly identical across training, validation, and test partitions, both for the original and new acquisitions (**Fig. B4**). If the model's accuracy relied on spatial autocorrelation within specific image acquisitions (as cautioned by Kattenborn et al.), performance would degrade significantly when evaluated on new NAIP imagery where those acquisition-specific patterns are absent. Furthermore, if the model were memorizing specific training locations, baseline error would be systematically lower on the training partition than on validation or test partitions. Neither pattern is present; the model maintains consistent accuracy across both spatial partitions and temporal epochs

***Temporal distance.*** *We confirmed via linear regression that error does not increase meaningfully with the temporal gap between NAIP acquisitions (1–9 years). The separation interval explained less than 2.4% of the variance ($r^2$ <0.024) across all evaluated error metrics, demonstrating mathematically that reported accuracy does not reflect short-lived, acquisition-specific signals.*

The dominant source of inter-epoch error variation is phenological mismatch between lidar reference and NAIP acquisition conditions, not acquisition-specific image characteristics. When phenology is controlled, performance degradation is negligible, consistent across data partitions, and independent of temporal separation — indicating that reported accuracy reflects learned canopy structure relationships rather than memorized acquisition-specific patterns.

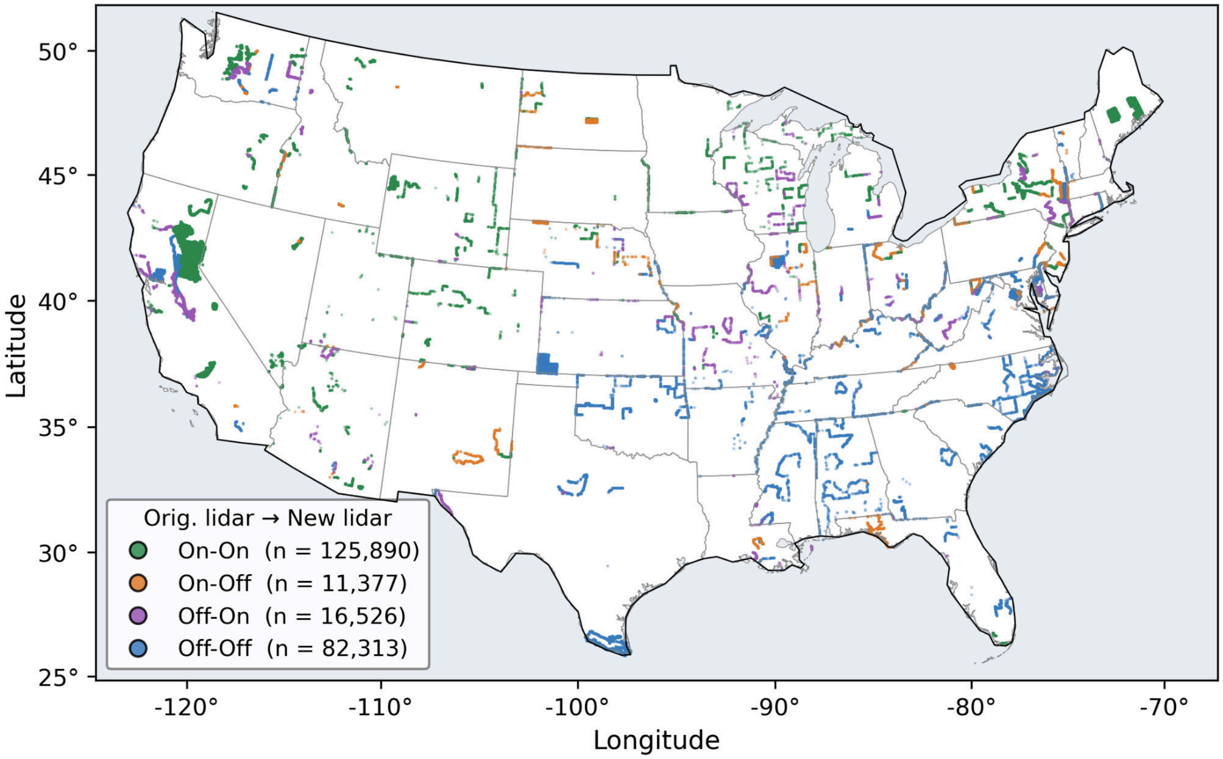


**Figure B1.** Geographic distribution of 228,123 matched tile locations, colored by leaf-status category (original lidar to new lidar).

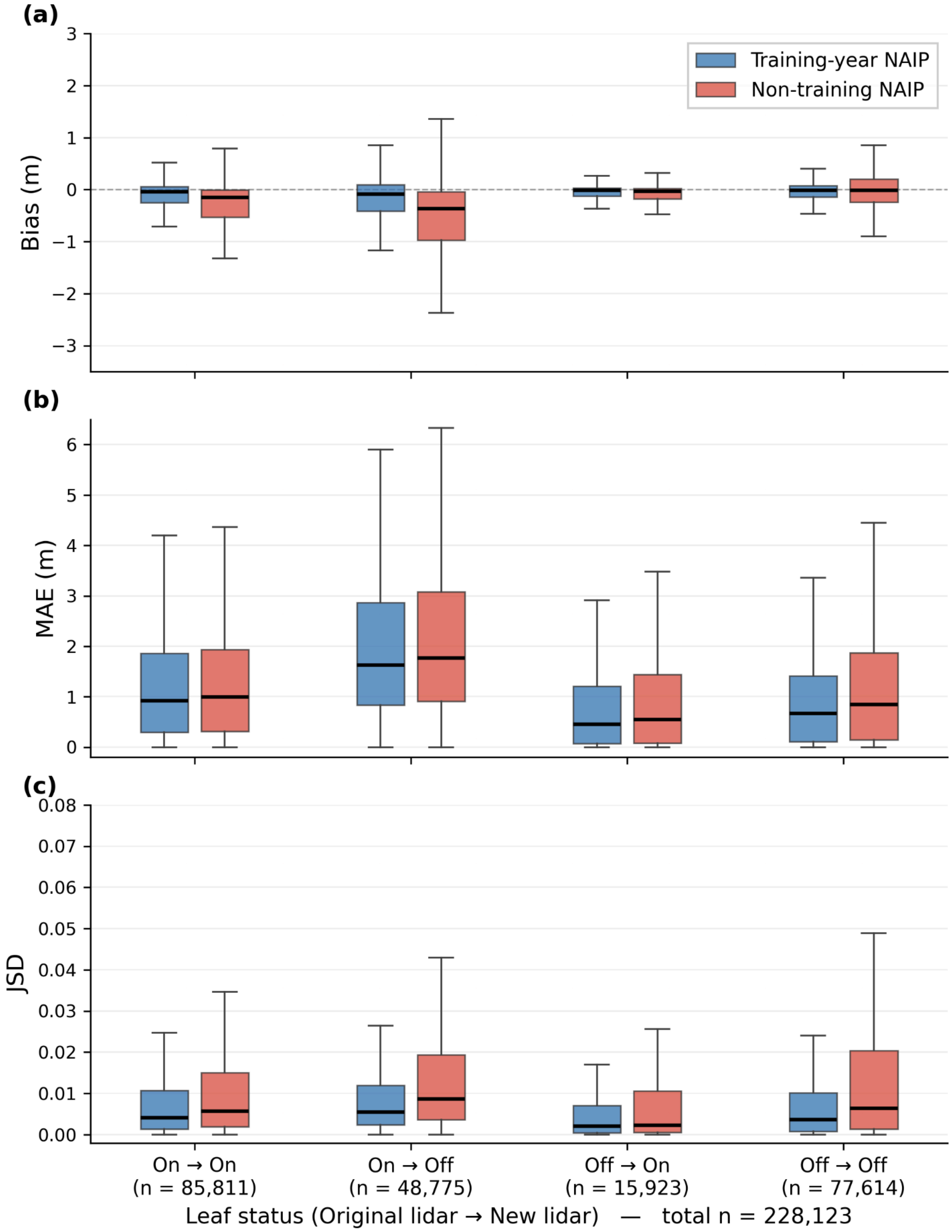


**Figure B2**: Paired boxplots of bias, MAE, and JSD for each leaf-status category, comparing training-year (blue) and non-training (red) NAIP predictions. The On-Off category shows the largest degradation; On-On shows minimal change.

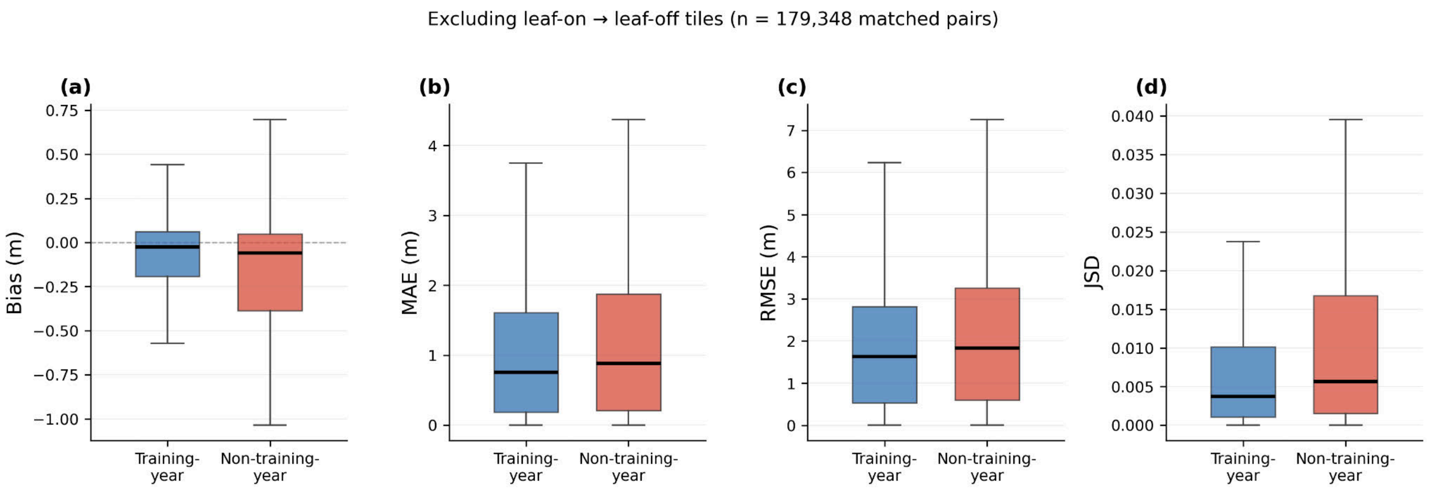


**Figure B3.** Paired boxplots of bias, MAE, RMSE, and JSD for training-year vs. non-training NAIP predictions, pooled across 179,348 matched tiles after excluding the On-Off phenological mismatch category.

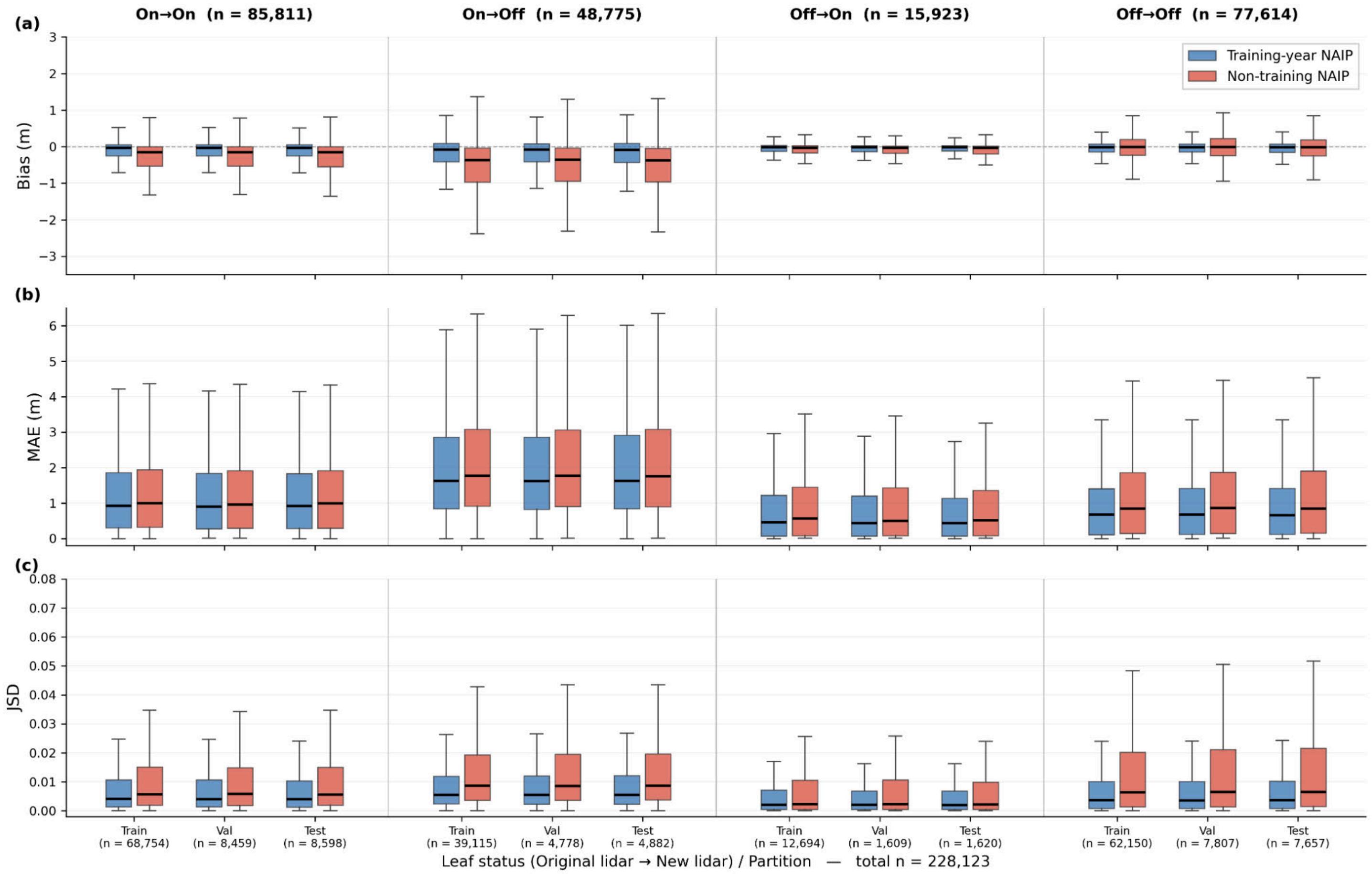


**Figure B4**: Same as Fig. B2, further stratified by data partition (train, validate, test). Error distributions are consistent across partitions within each leaf-status category.

#### Supplement C: Spatial Generalization

##### Overview & Methods

To assess whether the model generalizes to landscapes absent from training, we evaluated predictions across ten geographically isolated 10 × 10 km holdout regions distributed across the conterminous United States (Table C1, Figure C1) that were at a minimum 18 km from training data. Holdout sites were chosen to span a wide range of ecoregions and canopy structures — from arid shrubland and pinyon–juniper woodland in southern Nevada, to dry mixed conifer in eastern Washington, semi-arid rangeland in Montana, prairie and cropland in the Dakotas and Minnesota, mixed deciduous forest in the Appalachian piedmont (Virginia, Kentucky, New York), and dense coastal conifer forest on the southern Oregon coast.

Candidate regions were drawn from USGS 3DEP lidar projects[2] whose acquisition footprints fall entirely outside the geographic extent of the training data. To enforce strict spatial independence, we computed the Euclidean distance transform from all training-tile centroids on a 250 m EPSG:5070 (CONUS Albers) grid, and within each candidate project performed a 2 km-step grid search for the 10 × 10 km window that maximized the minimum distance from any training tile while remaining inside the project footprint. The resulting holdout sites lie 18–186 km from the nearest training tile (Table C1).

For each holdout region we generated an independent reference canopy height model from USGS 3DEP airborne lidar point clouds. Candidate lidar projects were selected from the USGS Work-unit Extent Spatial Metadata (WESM) catalog (*https://www.usgs.gov/3d-elevation-program/3dep-spatial-metadata*), restricted to recent vintage (2017 or later), and high quality level (QL 1 or QL 2). Reference CHMs were produced from the project LAZ tiles following the same procedure used to develop the training & validation dataset[6].

To place per-site accuracy in context, we constructed two reference distributions for each holdout region from the held-out test partition. The first is a geographic baseline: the 1,000 test tiles whose centroids lie closest (great-circle distance) to the holdout-region centroid, capturing the model's expected performance in the same general area. The second is a structural baseline: the 1,000 test tiles whose observed 95th-percentile canopy height most closely matches the holdout site's reference P95, capturing the model's expected performance on tiles with comparable canopy structure regardless of location. To make the holdout sites directly comparable to these tile-level distributions, we additionally split each 10 × 10 km holdout region into 256 x 256 m square sub-tiles (matching the typical footprint of a test tile) and computed per-sub-tile accuracy metrics. Sub-tiles with fewer than 50% valid pixels (after masking water, no data, and reference outliers) were excluded. The resulting three distributions per site — geographic neighbors, P95-matched tiles, and within-region sub-tiles — are compared in Figure C2. If the model generalized poorly to landscapes absent from training, we would expect to see substantial inflation in the holdout error distributions relative to both the geographic and structural reference sets.

##### Results

Table C2 presents per-site validation metrics computed across all valid pixels. Figure C2 contextualizes these results against two reference distributions for each holdout site... Across all ten regions, the model achieves an aggregate MAE of 1.66 m, RMSE of 2.81 m, $R^2$ of 0.80, and a mean JSD of 0.005. When excluding the single outlier region (NY Southeast) driven by reference data phenological mismatch, aggregate performance across the remaining nine regions improves to MAE = 1.31 m, RMSE = 2.30 m, and $R^2$ = 0.85. These values are consistent with the global and forest-specific statistics reported in the main manuscript.

As expected, absolute error tracks canopy height (Table C2). Sites dominated by open or sparse vegetation — sd_centralnorth (P95 = 0.80 m), montana (P95 = 7.82 m), nd_3dep (P95 = 10.18 m) — achieve MAEs at or below 0.35 m, while taller, denser sites (oregon P95 = 49.69 m; ny_southeast P95 = 24.35 m) carry larger absolute errors. Importantly, when each holdout site is compared against test tiles with similar canopy structure (Figure C2, green boxes), nine of the ten sites fall squarely within the expected error envelope. The Oregon site in particular, which initially appears to deviate from its geographic neighbors due to its much taller canopy (P95 49.7 m vs. 11.6 m for nearby tiles) produces error distributions that overlap closely with the P95-matched test tiles, confirming that the model's poorer performance at that location is a function of canopy height rather than a spatial generalization failure.

NY Southeast is a genuine outlier, which exhibits the largest errors of any holdout site (MAE = 4.83 m, bias = +3.21 m, $R^2$ = 0.30) and is the only error distribution that lies meaningfully outside both reference envelopes in Figure C2. Although the lidar and NAIP imagery were both acquired in 2022, they represent contrasting phenological states. The lidar was collected in late April; while this falls just inside the generalized "leaf-on" temporal heuristic used for continental-scale processing in Supplement B, the lidar data actually captured predominantly leaf-off conditions for the deciduous species in that area of New York – conditions that can persist into May. In contrast, the NAIP imagery was acquired in October during full leaf-on conditions. Because of this discrepancy, the model correctly predicts full-canopy height from the leaf-on NAIP imagery, but the leaf-off lidar reference captures a systematically reduced canopy surface—particularly for the deciduous broadleaf species that dominate the Appalachian/Hudson Valley landscape. The resulting positive bias is visible in the comparison chips (Fig. C3), where the model's canopy height predictions consistently exceed the sparse leaf-off reference. When evaluation is restricted to pixels with reference height > 5 m, the bias drops from +3.21 m to +1.58 m, showing that the overprediction is most pronounced where the model detects canopy foliage that the leaf-off reference fails to capture.

Together, these results support the conclusion that the UNetFiLM canopy height model generalizes well to geographic regions not represented in its training data. Across nine of ten holdout sites spanning arid shrubland, semi-arid rangeland, prairie, mixed-conifer, eastern deciduous forest, and dense coastal conifer forest, the model achieves accuracy comparable to test-set tiles drawn from both nearby geography and structurally similar canopies elsewhere in CONUS, and we do not observe systematic inflation of errors. The single deviation, NY Southeast, is fully attributable to a phenological mismatch between leaf-off reference lidar and leaf-on input imagery, not to spatial generalization failure. These findings complement the temporal acquisition sensitivity analysis and are consistent with the finding that the model has learned generalizable canopy-structure relationships rather than spatially or temporally specific image patterns.

**Table C1**: Holdout Regions

| Site Name | USGS Lidar Project* | Lidar Date | NAIP Date | Distance to Training Data (km) | Vegetation |
|---|---|---|---|---|---|
| sd_centralnorth | SD_CentralNorth_D22 | 11/11/23 | 10/19/23 | 29 | Cropland, prairie, very sparse canopy |
| montana | MT_Statewide_Phase5_D23 | 8/24/23 | 7/21/23 | 29 | Semi-arid rangeland, conifer woodlands |
| nd_3dep | ND_3DEPProcessing_D22 | 5/9/17 | 10/7/17 | 186 | Cropland, riparian corridors |
| mn_becker | MN_BeckerCounty_2021_D21 | 5/15/22 | 7/16/21 | 67 | Mixed deciduous/conifer, lakes, wetland |
| nevada | NV_Southern_D23 | 10/31/23 | 6/17/22 | 47 | Arid shrubland and pinyon-juniper |
| wa_northeast | WA_NorthEast_B22 | 4/6/22 | 6/30/23 | 18 | Dry mixed conifer |
| va_southcentral | VA_FEMA_NRCS_SouthCentral_2017_D17 | 4/14/17 | 8/22/18 | 24 | Mixed deciduous/conifer |
| ky_centraleast | KY_CentralEast_A23 | 12/12/22 | 5/30/22 | 58 | Mixed deciduous forest |
| oregon | OR_SouthCoast_2019_A19 | 10/17/20 | 6/17/20 | 91 | Dense conifer forest |
| ny_southeast | NY_SouthEast4County_A22 | 4/24/22 | 10/21/22 | 25 | Deciduous forest |

* Lidar data downloaded from https://rockyweb.usgs.gov/vdelivery/Datasets/Staged/Elevation/LPC/Projects/

**Table C2**: Per-site validation metrics computed over all valid pixels. Reference height summaries (mean and 95th percentile, in m) characterize the canopy structure of each holdout site.

| Region | pixels | Ref. mean (m) | Ref. P95 (m) | Bias | MAE | r2 | RMSE | JSD |
|---|---|---|---|---|---|---|---|---|
| sd_centralnorth | 98,612,003 | 0.17 | 0.8 | -0.074 | 0.112 | 0.800 | 0.411 | 0.009 |
| montana | 96,697,432 | 0.97 | 7.82 | -0.058 | 0.309 | 0.884 | 0.888 | 0.0004 |
| nd_3dep | 97,876,740 | 1.06 | 10.18 | 0.088 | 0.350 | 0.863 | 1.378 | 0.0012 |
| mn_becker | 61,707,679 | 3.55 | 17.58 | -0.145 | 0.520 | 0.964 | 1.192 | 0.0012 |
| nevada | 93,693,042 | 2.43 | 6.8 | -0.319 | 0.752 | 0.758 | 1.174 | 0.007 |
| wa_northeast | 88,459,915 | 4.44 | 19.58 | 0.015 | 1.295 | 0.861 | 2.532 | 0.0013 |
| va_southcentral | 91,214,148 | 11.33 | 25.94 | -0.349 | 1.701 | 0.887 | 3.051 | 0.0013 |
| ky_centraleast | 95,505,137 | 11.97 | 28.2 | -0.087 | 2.104 | 0.883 | 3.563 | 0.0047 |
| oregon | 48,407,806 | 24.09 | 49.69 | -0.956 | 4.641 | 0.785 | 6.519 | 0.0075 |
| ny_southeast | 81,176,737 | 9.19 | 24.35 | 3.210 | 4.830 | 0.301 | 7.407 | 0.0274 |

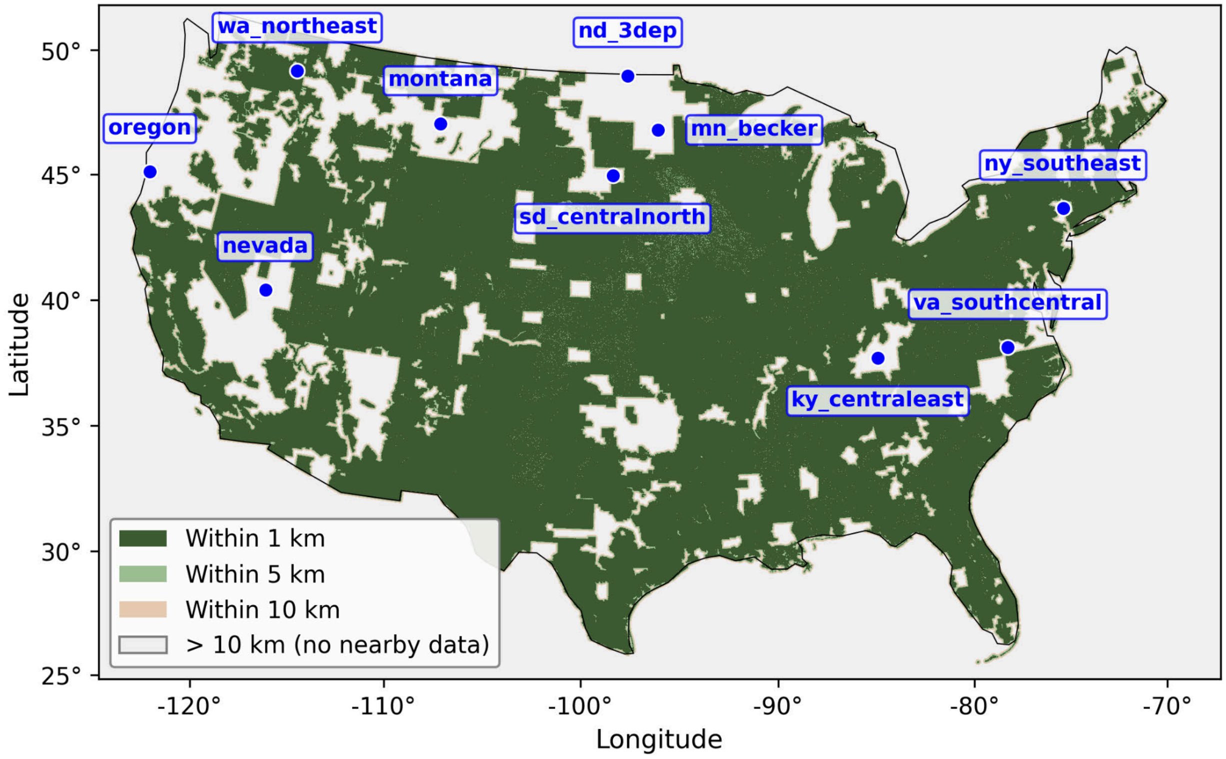


**Figure C1:** Locations of holdout samples used in this analysis. The Basemap shows the coverage of the CHM & NAIP chips used in our modeling exercise, colored by maximum distance to a training chip for each pixel.

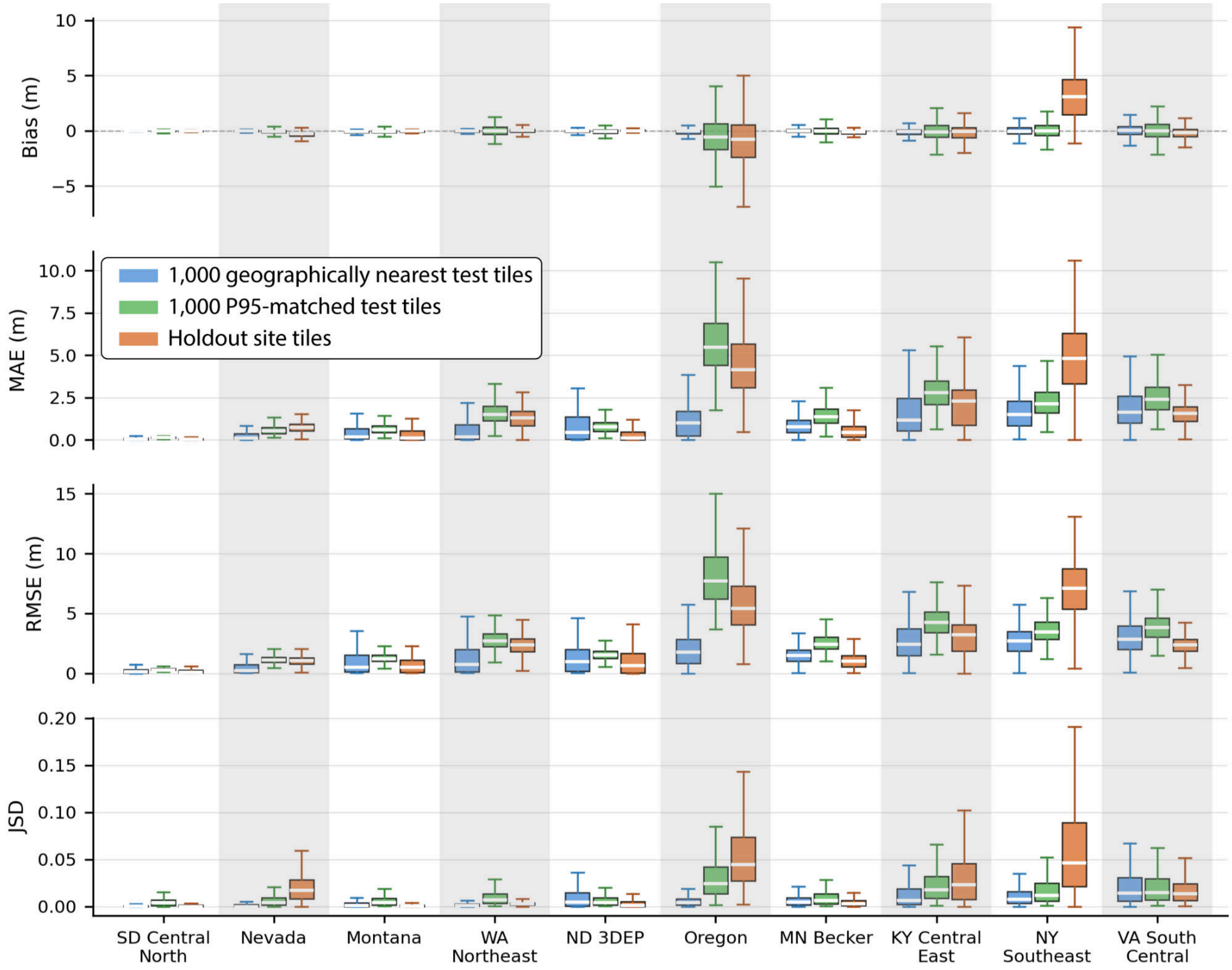


**Figure C2:** Validation metrics (bias, MAE, RMSE, and JSD) for ten holdout sites, computed across ~256 m sub-tiles (orange boxplots). To contextualize performance, holdout metrics are compared against two reference distributions drawn from the global test data: the 1,000 geographically nearest test tiles (blue boxplots) and the 1,000 structurally similar test tiles matched by 95th-percentile (P95) canopy height (green boxplots). Sites are ordered by increasing mean canopy height of nearby tiles. Holdout metrics generally align with expected performance based on the global data and surrounding regional baselines, with two exceptions: Oregon, where elevated errors result from an exceptionally tall canopy relative to geographic neighbors (though in line with structurally similar global tiles), and NY_southeast, where errors are driven by a leaf-on/leaf-off phenological mismatch between the NAIP imagery and lidar reference.

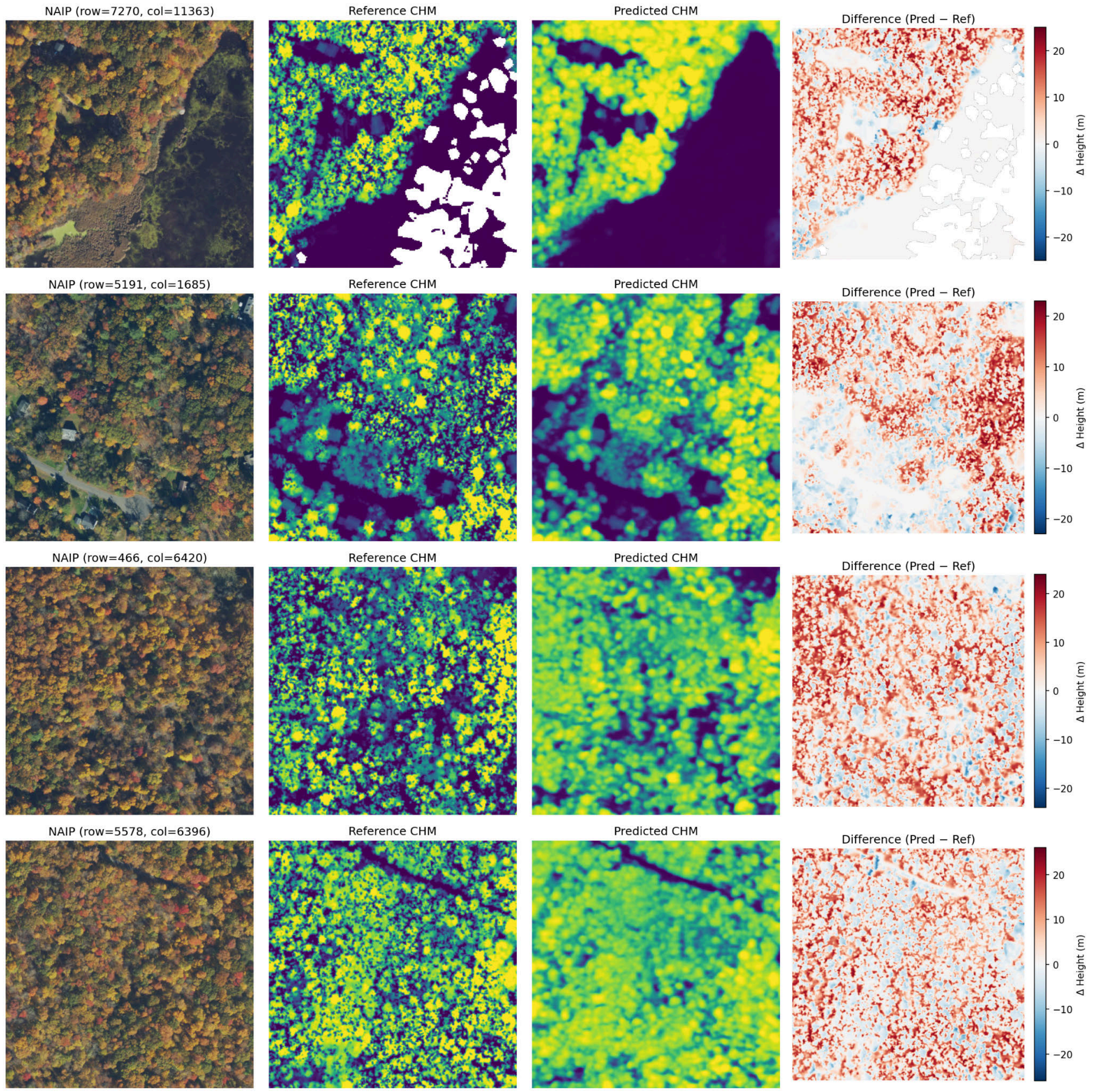


**Figure C3**. Representative chip comparisons for the NY_southeast site, where a leaf-season mismatch between the October 2022 NAIP (leaf-on) and April 2022 lidar reference (leaf-off) drives systematic overprediction. Columns show NAIP RGB, reference CHM, predicted CHM (shared color scale), and the prediction residual (Pred − Ref); warm colors in the difference column indicate positive bias.
